# Accurate estimation of cell-type resolution transcriptome in bulk tissue through matrix completion

**DOI:** 10.1101/2021.06.30.450493

**Authors:** Weixu Wang, Xiaolan Zhou, Jun Yao, Haimei Wen, Yi Wang, Mingwan Sun, Chao Zhang, Wei Tao, Jiahua Zou, Ting Ni

**Affiliations:** State Key Laboratory of Genetic Engineering, Collaborative Innovation Center of Genetics and Development, Human Phenome Institute, School of Life Sciences and Huashan Hospital, Fudan University, Shanghai, 200438, P.R. China; Ministry of Education (MOE) Key Laboratory of Contemporary Anthropology, Human Phenome Institute, School of Life Sciences, Fudan University, Shanghai, 200438, P.R. China; MOE Key Laboratory of Cell Proliferation and Differentiation, School of Life Sciences, Peking University, Beijing, 100871, P.R. China; Key Laboratory of Gene Engineering of the Ministry of Education and State Key Laboratory of Biocontrol, School of Life Sciences, Sun Yat-Sen University, Guangzhou, 510006, P.R. China; Guangdong Provincial Key Laboratory of Bioengineering Medicine, National Engineering Research Center of Genetic Medicine, Institute of Biomedicine, College of Life Science and Technology, Jinan University, Guangzhou, 510632, P.R. China

## Abstract

Single cell RNA-seq (scRNA-seq) has been widely used to uncover cellular heterogeneity, however, the constraints of cost make it impractical as a routine on large patient cohorts. Here we present ENIGMA, a method that accurately deconvolute bulk tissue RNA-seq into single cell-type resolution given the knowledge gained from scRNA-seq. ENIGMA applies a matrix completion strategy to minimize the distance between mixture transcriptome and weighted combination of cell type-specific expression, allowing quantification of cell type proportions and reconstruction of cell type-specific transcriptome. The superior performance of ENIGMA was validated in simulated and realistic datasets, including disease-related tissues, demonstrating its ability in novel biological findings.

## Background

Each cell type has unique role in the ecosystem of tissues. In recent 20 years, from microarray to Next Generation Sequencing (NGS), gene expression quantification has been a useful resource for understanding the complex regulation related to different diseases or phenotypes[1, 2]. There are a lot of large cohort studies, such as TCGA[3] and GTEx[4], to build the connection between gene expression and various phenotypes. The sequenced samples are the collection of various cell types (bulk tissue), therefore the expression profile of a given sample is the convolution of gene expression from heterogenous cell types. It has been shown that cell type-specific gene expression alteration can contribute to our understanding of the principles of biological regulation[5]. Therefore, it is necessary to isolate cell type-specific gene expression profile from mixed bulk tissue.

To this end, researchers have pursued several advanced techniques such as fluorescence-activated cell sorting (FACS)[6] and single-cell RNA-sequencing (scRNA-seq)[7] to quantify cell type-specific expression (CSE) profile at sample level. These technologies bring new insights to us about the landscape within the tissues, but they still have some limitations. For FACS dataset, it relies on small combinations of preselected marker genes, limiting the cell types that could be studied. For scRNA-seq dataset, although it could collect a large number of cells from tissues, and characterize a much richer heterogenous cell state information compared with FACS, its relatively high cost property does not allow the routine assessment of tissues from hundreds of samples, thus limits its application on large cohort cell type-specific phenotype-gene expression association analysis[8].

Previous researches have pursued to integrate FACS and single-cell RNA-seq datasets to help us deconvolute the bulk RNA-seq dataset. The methods such as CIBERSORT[9], TIMER[10], BSEQ-sc[11] and MuSiC[12], used several regression models to deconvolute the bulk RNA-seq dataset into cell type fraction matrix, and helped us to infer the proportion information of each cell type. Although these methods have been widely applied in previous studies to help find cell type fractional variation, they could not provide the cell type-specific expression profile at sample level. Besides, there were methods that had been developed for deconvolving cell type-specific gene expression at sample level (e.g. bMIND[13] and CIBERSORTx[9]). bMIND uses full Bayesian model, which bases on cell type reference matrix derived from single cell RNA-seq as the prior information, and applies Markov Chain Monte Carlo (MCMC) to estimate parameters[13]. As bMIND estimates the cell type-specific expression for each gene independently, it ignores the nature of latent correlation among genes, which is important for gene co-expression network analysis[14]. CIBERSORTx estimate CSE profile based on non-negative least squares algorithm with the goal of separating two groups of samples[8], but it cannot perform deconvolution for all inputted genes and cell types. In addition, these two methods are all time-consuming, and the expensive time consumption makes the parameter tuning and optimization not very convenient.

To address these deficiencies, we developed a d<u>e</u>co<u>n</u>volut<u>i</u>on method based on regularized <u>ma</u>trix completion (ENIGMA) to refine the estimation of CSE profile for each bulk sample in an extremely short time. To make the inferred CSE reliable, we minimized the distance between each inferred CSE average expression level and the reference expression profile. We validated its inference accuracy on both simulated and real datasets through showing its superior performance compared with other state-of-the-art methods. We proved that ENIGMA could give better CSE estimation as reflected by its inferred gene level or sample level correlation with simulated ground truth CSE under different expression noise and reference noise setting. Also, it showed better cell type-specific differential expression gene (DEG) identification compared with other algorithms. Moreover, we applied ENIGMA on three different scenarios, and proved that it could identify unique cell type-specific expression pattern and co-expression network. We identified the differentiation trajectory of monocytes in arthritis patients. Meanwhile, we found a senescence-associated gene co-expression module in pancreas islet beta cells, which is reported relevant to type 2 diabetes (T2D) patients.

## Results

### The overview of ENIGMA

We illustrated the general strategy of ENIGMA about performing bulk gene expression deconvolution to infer CSE at sample level. We regarded this task as a matrix completion problem, with bulk gene expression matrix could be represented as the linear combination of several cell type-specific expression matrices. The linear combination coefficient is cell type fraction (*θ_i_*), which could be estimated through using a cell type fraction estimation algorithm or experimental measurements through cell sorting. Our goal is to minimize the distance between observed bulk gene expression matrix and reconstituted bulk expression matrix through linear combination of inferred CSE (Fig. 1a, b).

**Fig. 1:**
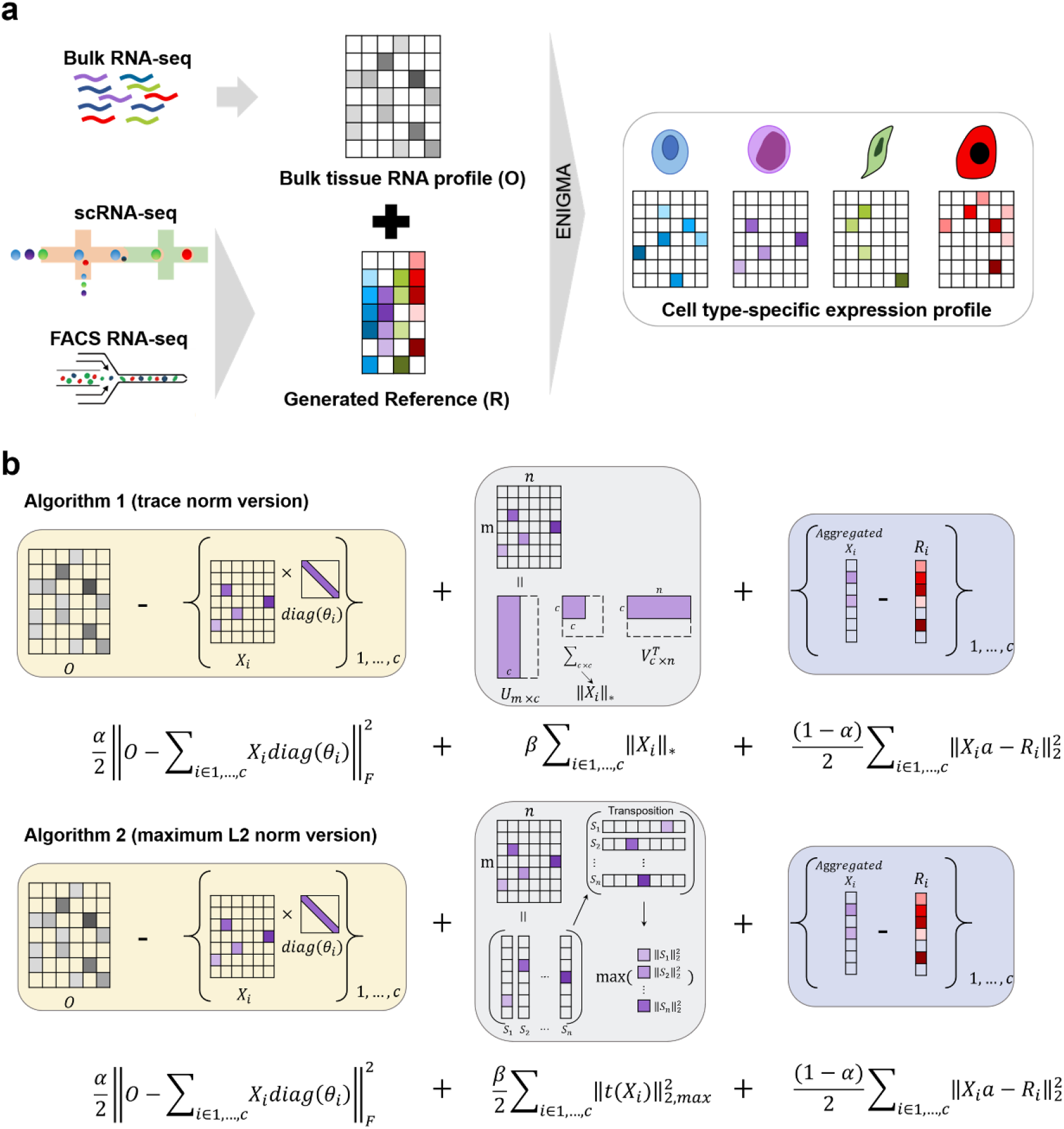
Cell type-specific expression (CSE) inference using ENIGMA. a. Overview of the ENIGMA algorithm. ENIGMA aims to integrate cross-platform RNA-seq datasets to infer the cell type-specific gene expression (CSE) profile. ENIGMA borrows the knowledges from single cell RNA-seq and FACS RNA-seq information to generate reference profile to aid CSE profiles inference. b. Schematic overview of the ENIGMA algorithm. The loss (object) function has three terms. The first term means the difference between the observed bulk gene expression matrix and the linear combination of the CSE profiles. The mixing coefficient for each CSE is quantified as the cell fraction matrix, which could be estimated from classical cell type deconvolution algorithms (e.g. CIBERSORT, Robust Linear Regression). The second term is regularization term, which can be used to control the rank of CSE or shrink the CSE (extreme values in a CSE are “shrunk” towards a central value). The third term represents the difference between aggregated inferred CSE and reference profile across each cell type.

Besides the minimization of the main object function, we also added two additional constraint terms to make the inference more robust. First, we considered each aggregated CSE should be as close as to the corresponding reference, which could also be regarded as the prior expression information derived from reference matrix (Fig.1b). Second, we used regularization term to constrain the trace norm (sum of singular values) or maximum ℓ_2_ norm on inferred CSE (Fig. 1b). Both constraints were from two fundamental hypotheses of our CSE deconvolution models (Method). The numerical analyses also supported that using these two regularization terms would benefit the CSE profile estimation (see Loss design section of **Supplementary Notes**).

ENIGMA has three main steps (Figure S1). First, ENIGMA requires cell type reference expression matrix (profile matrix), which could be derived from either FACS RNA-seq or scRNA-seq datasets through calculating the average expression value of each gene from each cell type. Previous researches have widely used reference matrix curated from different platforms, for instance, Newman et al. used LM22 immune signature matrix which derived from microarray platform to deconvolute bulk RNA-seq dataset[9]. However, we have to note that using references from different platforms would introduce unwanted batch effect between reference and bulk RNA-seq matrix, especially for the reference matrix derived from low coverage scRNA-seq dataset. To overcome this challenge, we used a previously presented method that was specifically designed for correcting batch effect among bulk RNA-seq matrix and reference matrix (Method)[8]. Second, ENIGMA applied robust linear regression model[15] to estimate fractions of each cell type among samples based on reference matrix derived from the first step. Third, based on reference matrix and cell type fraction matrix estimated from step 1 and step 2, ENIGMA applied constrained matrix completion algorithm to deconvolute bulk RNA-seq matrix into CSE profiles at sample level. In order to constrain the model complexity to prevent overfitting, we applied two different norm penalty functions to regularize resulted CSE. Finally, the returned CSE could be used to identify cell type-specific DEG, visualize each gene’s expression pattern on the cell type-specific manifold space (e.g. t-SNE[16], UMAP[17]), and build the cell type-specific co-expression network to identify modules that are relevant to phenotypes of interest.

### ENIGMA gives better estimation of CSE profile at sample level

We benchmarked the performance of ENIGMA with other CSE inference tools with simulated datasets and realistic datasets. First, we tested whether ENIGMA could yield a better gene expression inference compared with other algorithms. For comparison, we employed two other methods: 1) TCA (Tensor Composition Analysis), a frequentist approach that is designed for bulk DNA methylation data and can also be applicable for CSE estimation[18]; 2) bMIND, a full Bayesian model that models the gene expression as Gaussian distribution with the prior information from reference matrix and uses MCMC to learn the parameters[13]. We compared the performance of two versions of ENIGMA, which correspond to their norm functions for regularization (see Method for details), to these two algorithms. An ideal CSE inference algorithm should recover the cell type-specific sample variation and gene variation. We tested this by generating simulated bulk RNA-seq dataset through linear combination of the simulated cell type-specific expression profiles generated from single cell RNA-seq data (see **Supplementary Notes** for details). After reconstructing CSE based on each algorithm, we calculated sample level correlation and gene level correlation between inferred CSE and ground truth CSE in each simulated cell type.

We benchmarked the CSE inference on pseudo bulk RNA-seq datasets generating from scRNA-seq of head and neck squamous cell carcinoma (HNSCC)[19] and melanoma scRNA-seq dataset[20] (see Methods for details). The results indicated that ENIGMA performed either comparable or better than TCA and bMIND on seven cell types regardless sample level or gene level correlation coefficients in HNSCC (Fig. 2a) or melanoma patients (Fig. 2b). We also compared gene level correlation after performing quantile normalization to each inferred CSE, and both versions of ENIGMA still performed consistently better than TCA and bMIND (Figure S2a). We then extended our analysis to the real bulk RNA-seq dataset with corresponding FACS RNA-seq as the ground truth. A previous study used surgically resected primary non-small cell lung cancer (NSCLC) tumor biopsies (n = 24 patients) to generate RNA-seq libraries of four major subpopulations purified by FACS: epithelial/cancer, immune, endothelial and fibroblast cells [21], which were served as the ground truth for further analysis. After deriving a reference matrix from external NSCLC scRNA-seq datasets (Method) [21], we applied ENIGMA to perform *in-silico* expression purification on the bulk samples, and compared ENIGMA with TCA, bMIND and additional algorithms, which merely deconvolute limited genes or cell types (CIBERSORTx[8], ISOpure[22] and Demix[23]). ENIGMA had higher Spearman correlation between inferred CSE and ground truth (FACS-purified tumor cells) CSE per sample than all other methods (Fig. 2c upper panel). Even though ENIGMA had lower Spearman correlation at gene level compared with TCA (Fig. 2c lower panel), we observed that ENIGMA gave better gene level estimation on other cell types including fibroblasts and endothelial cells (Figure. S2b). Overall, compared with previous algorithms, ENIGMA gave a more accurate estimation of CSE profiles.

**Fig. 2:**
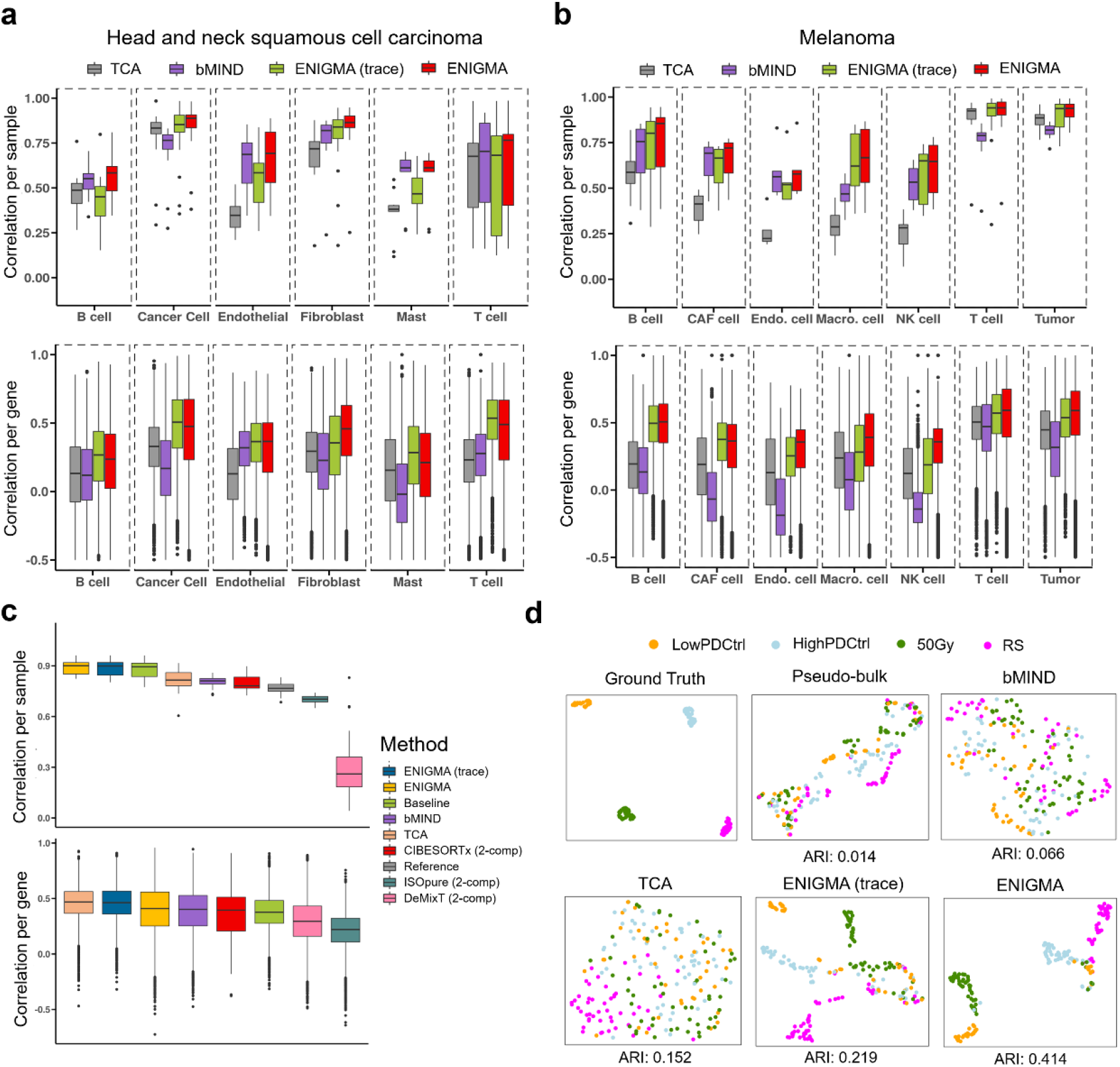
Validation of ENIGMA on simulation data. a. Each imputed CSE profile was compared with its corresponding ground truth CSE by Spearman correlation. For each cell type, we computed correlation across genes for each sample (upper panel) and correlation across samples for each gene (lower panel). The ground truth CSE and bulk expression profile for deconvolution was generated from scRNA-seq dataset of head and neck squamous cell carcinoma. b. Each imputed CSE profile was compared with its corresponding ground truth CSE by Spearman correlation. For each cell type, we computed correlation across genes for each sample (upper panel) and correlation across samples for each gene (lower panel). The ground truth CSE and bulk expression profile for deconvolution was generated from scRNA-seq dataset of melanoma. c. The imputed NSCLC tumor specific profile was compared with its corresponding ground truth CSE by Spearman correlation. We computed correlation across genes for each sample (upper panel) and correlation across sample for each gene (lower panel). The ground truth CSE and bulk expression profile for deconvolution was curated from previously sequenced dataset of NSCLC. The Spearman correlation coefficient (Y-axis) was computed based on gene set filtered by CIBERSORTx. The *P* value was calculated from paired Wilcoxon rank sum test. d. The Uniform Manifold Approximation and Projection (UMAP) plots of gene expression profiles. Each dot represented a simulated sample and colored according to the fibroblasts senescence labels (LowPDctrl (young quiescent cells), HighPDctrl (middle quiescent cells), 50Gy (X-ray induced senescent cells), RS (replicative senescent cells))[24]. The simulated ground truth CSE profile was generated from scRNA-seq dataset of senescent fibroblasts (left of the upper panel), labeled as “Ground Truth”. The middle of the upper panel denoted ground truth profile mixed with other two simulated CSE profiles to generate a simulated bulk RNA expression profile, labeled as “Bulk”. We further used bMIND, TCA, ENIGMA and ENIGMA (trace) to deconvolute the admixture sample to infer the fibroblasts expression profile, and inspected whether the algorithms recover the latent cell clustering structure through visualization on UMAP plots. The clustering performance was indicated through Adjusted Rand Index (ARI).

To further support the robustness of our algorithm on estimating CSE profiles, we simulated pseudo-bulk with four different levels of observation noise (0.1, 0.5, 1, 1.5). The correlation per sample and per gene between ENIGMA inferred CSE and ground truth CSE was comparable to those from TCA and bMIND at low-noise levels and was higher than those from TCA and bMIND at high-noise levels (Figure S2c). It also showed that the performance of maximum ℓ_2_ norm-based regularization (default) ENIGMA was better than trace norm-based regularization ENIGMA at different noise levels (Figure S2c).

### ENIGMA recovers latent cell states for each cell type

In CSE inference, recovering the correlation structure among samples is also a very important aspect, which could help us to identify the latent higher-order data structure in sequenced cohort at cell type level. However, this important aspect was ignored by previous CSE inference strategies[13, 18]. To benchmark this aspect, we firstly used scRNA-seq dataset sequenced from fibroblasts with four different senescence states [24], combining with two simulated scRNA-seq datasets [25] to generate a simulated ground truth CSE profile. Next, we mixed the simulated fibroblasts CSE with other two simulated expression profiles representing two different cell types with specific proportion to constitute a pseudo-bulk expression profile (Figure S3a, also see Benchmark tasks and metrics section of Supplementary Notes). We found that using any existing data processing methods would not help to recover the latent cell states of fibroblasts (Figure S3b). After deconvolution, the result showed that both versions of ENIGMA (maximum ℓ_2_ norm and trace norm) could recover the four distinct senescence states of simulated fibroblast CSE with relatively high accuracy (Fig. 2d, Figure S3c, ARI = 0.219 for ENIGMA (trace); ARI = 0.414 for ENIGMA), much better than other methods. Even though we observed that some of the samples were not grouped into their corresponding cell clusters (marked with dashed circle in Figure S3d), those samples had low abundance of fibroblasts (Figure S3d) and previous research had proved that deconvolution on low abundant cell types would gain bad estimation of CSE[18].

To explore whether deconvolution methods could reconstruct the latent continuous structure of CSE at sample level, we simulated the CSE which has latent sample-wise correlation structure that could be organized as a transition trajectory on the diffusion component space, and attached each sample a pseudo-time label to show the relative order information[26]. Then, we mixed this CSE profile with other two simulated CSE matrices representing three different cell types with specific proportion to build a pseudo-bulk expression profile. After deconvolution, the result showed that only ENIGMA (maximum ℓ_2_ norm) could recover the trajectory structure of the specific CSE, while ENIGMA (trace), TCA and bMIND totally lost this information (Figure S3d). Taken together, these results demonstrate that ENIGMA could infer CSE with better estimation at sample level concordance, and could recover the correlation structure of CSE profiles better than previous methods.

### The robustness and scalability of ENIGMA

As reference matrix is extremely important for CSE inference and different sources of reference matrices (library type, sequence depth, batches, etc.) may introduce variability of the reference, we therefore evaluated the influence of the quality of reference matrix by adding variable noises at different levels in the simulated data. We found that although sample level correlation between inferred and ground truth CSE was not distinctly affected, the gene level correlation was clearly influenced by the noise of reference matrix (Figure. S4a), especially for TCA and bMIND. The maximum gene level correlation coefficient decrease could reach 0.26 in TCA and 0.32 in bMIND, while the maximum correlation coefficient decreases just reach 0.2 and 0.06 in ENIGMA and ENIGMA (trace), respectively (Figure. S4b). We noticed that the quality of reference matrix also seriously influenced the prediction accuracy of cell type fraction matrix, with the average correlation coefficient drops from 0.94 to 0.4 (Figure. S4c). We also compared the all-gene level correlation coefficient decrease among four methods, and found that both ENIGMA and ENIGMA (trace) showed significantly better robustness compared with TCA and bMIND (*P* value < 2.2e-16, Figure. S4d).

Next, we tested the influence of the number of latent cell types in bulk gene expression matrix to the CSE inference through admixing a range of cell type numbers to build bulk gene expression matrices. The results indicated that both versions of ENIGMA performed consistently better than TCA and bMIND regardless the number of cell types we simulated (Figure. S5).

Efficiency of computing is very important for CSE inference since faster runtime will greatly facilitate parameter tuning and optimization especially analyzing large cohort datasets. To evaluate ENIGMA’s computational performance, we measured total runtime usage of ENIGMA versus other methods. We created five benchmark datasets with 100, 500, 1000, 5000, and 10000 genes with fixed sample size (100 samples), and reported the runtime for each condition. The results showed that ENIGMA’s runtime scaled well for all datasets (ranging from 4 to 24 seconds), while runtime of the other two methods (TCA and bMIND) showed exponential increase when gene numbers rose from 100 to 10000 (Figure. S6). These results demonstrate that ENIGMA is computationally efficient and capable of analyzing all genes with large sample size on servers or even personal computers.

### ENIGMA benefits cell type-specific-DEGs (CTS-DEGs) identification

A well inferred CSE also could help us to identify cell type-specific differentially expressed genes (CTS-DEGs) associated with cohort phenotypes[8, 13]. To evaluate ENIGMA’s ability to discover the CTS-DEGs, we generated simulated dataset with known DEGs for each cell type, and calculated the sensitivity and specificity for the identified CTS-DEGs. Besides TCA and bMIND, we also ran differential expression analysis on raw simulated bulk dataset as baseline for comparison.

We applied similar benchmark evaluation as Rahmani et al[18], and assessed sensitivity and specificity as the function of effect size (the magnitude of expression differences). For each CSE profile, we applied ordinary least square to fit the regression model for each inferred gene expression and simulated phenotype in each cell type, and defined those genes with False Discovery Rate (FDR) adjusted *P* value < 0.05 as the differentially expressed genes (DEGs). We found that ENIGMA showed better sensitivity at low effect size compared with TCA and bMIND (Fig. 3a, b). Meanwhile, ENIGMA (trace) showed consistently higher sensitivity compared with other algorithms in all scenarios (Fig. 3a, b), indicating that ENIGMA tends to identify more true CTS-DEGs compared with other algorithms. To reveal that ENIGMA could refine the phenotype-associated expression difference, we compared the alteration (expression difference) we estimated from mixed expression profile and ENIGMA inferred CSE with the ground truth in each cell type. We found that the correlation between gene expression alteration estimated from ENIGMA inferred CSE and ground truth was better than the correlation between gene expression alteration estimated from mixed expression profile and ground truth (Fig. 3c, d), supporting that ENIGMA could find the cell type-specific phenotype associated alterations that diminished or enwombed in mixed expression profile by the sample-wise mixing proportions of different cell types. We also performed power analysis through assessing the influence of sample size to the sensitivity and specificity of CTS-DEGs identification, and reflected both of them using the Area Under Curve (AUC) value. The results showed that ENIGMA (trace) and bMIND tended to perform consistently better than other algorithms across different sizes of samples (Figure S7). Moreover, ENIGMA (trace) showed better performance (higher AUC) than other methods when effect size increased (Figure S7).

**Fig. 3:**
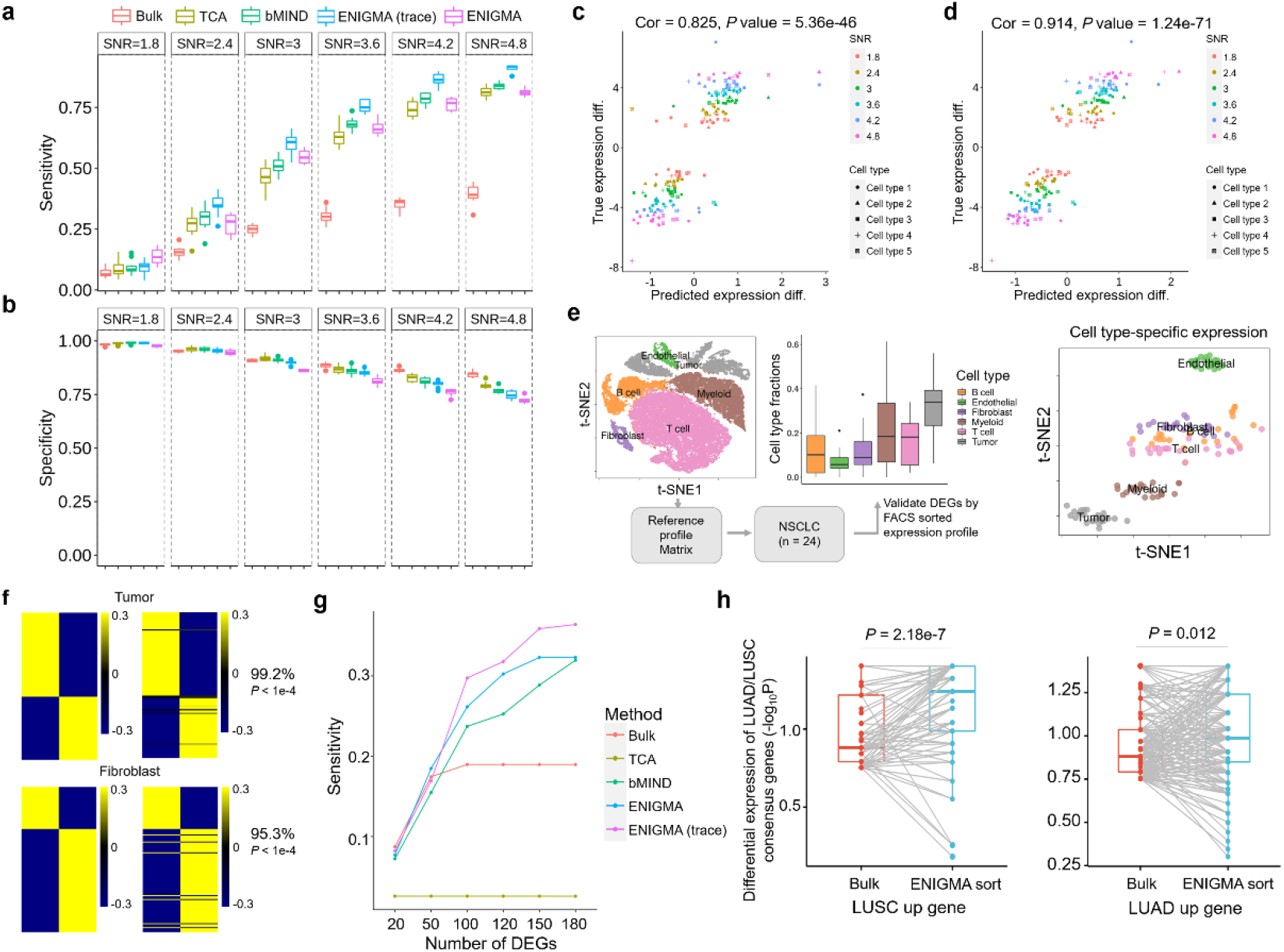
Benchmark of ENIGMA on CTS-DEGs detection. a-b. Boxplots of sensitivity (a) and specificity (b) for the evaluation of detecting cell type-specific differentially expressed genes (CTS-DEGs) on the simulation dataset. The rate of true positives (sensitivity) and the rate of true negatives (specificity) were measured under a range of signal to noise ratio (SNR). All of the simulations were evaluated under the setting of five constituting cell types, 1000 genes, 100 samples and Gaussian observation noise with unit variance. Each cell type had 100 DEGs with randomly down- or up-regulated. c,d. Scatterplots of the true gene expression difference (diff.) in affected cell types (Y-axis) versus predicted gene expression difference (X-axis) from raw bulk RNA profile (c) or CSE profile imputed from trace-norm based ENIGMA (d). Data points were shown for 500 DEGs, five affected cell types and six SNR levels. Each point was colored according to SNR level and shaped by the cell type it comes from. e. Schema of deconvoluting NSCLC tumor tissue bulk RNA profile using ENIGMA. The reference matrix was constructed from previous scRNA-seq dataset, and it was applied on the bulk RNA profile from 24 patients to deconvolute the bulk RNA profile into cell type fraction matrix and cell type-specific expression profile. Left: A t-SNE plot showing eight major tumor subpopulations profiled from six donors by scRNA-seq. Each dot represents a cell and is colored according to the cell type. Middle: A barplot showing eight major tumor subpopulation fractions in 24 patients[27]. Right: A t-SNE plot showing eight major tumor subpopulations deconvoluted from bulk RNA profile of 24 patients. Each dot represents a patient and is colored according to the cell type it belongs to. f. Analysis of the concordance between ENIGMA-imputed (left) and flow-sorted (right) CTS-DEGs expression profiles in LUAD and LUSC. The heatmaps show the averaged mean-centered log2-adjusted expression values for each DEG identified by ENIGMA. The DEGs were defined as the genes with |log2FC| > 2 and adjusted *P* value < 0.01. ‘DEG concordance’ denotes the fraction of genes with the same direction of differential expression between ENIGMA and flow-sorted profile. The *P* value was calculated through Monte Carlo strategy (Method). g. The curves showing the sensitivity vs. the number of selected top DEGs. The sensitivity was calculated based on the number of top DEGs estimated from each method. The top DEGs (adjusted *P* value < 0.05) were ranked through |log2FC|. For each method, when the number of top DEGs exceeded the total number of the DEGs, the sensitivity was calculated according to the total DEGs. The pseudo ground truth DEGs were defined from flow-sorted profile and scRNA-seq profile (Method). h. Significance comparison between bulk RNA profile and ENIGMA-imputed profile for pseudo ground truth DEGs from panel g. The significance (-log10(*P* value)) was calculated by Wilcoxon rank-sum test.

Besides sensitivity and specificity, the precision, or 1-false discovery rate, is also very important metric for DEGs identification. But using ordinary least regression model would lead to the high false discovery rate of DEGs identification. In order to fairly benchmark different methods, we used an FDR controlled DEGs calling method to perform CTS-DEGs analysis. We found that ENIGMA (trace) performed consistently better than other methods on sensitivity (Figure S8a), but along with relatively higher false discovery rate (Figure S8b). However, if we controlled the number of called DEGs by ENIGMA (trace) as the same number as other method (TCA), the FDR of ENIGMA (trace) was comparable or even lower than other methods (Figure S8b). It suggested that the criteria (the threshold for *P* value) for calling DEGs would influence the benchmarks, so we plotted the precision-recall curve (precision is equivalent to 1-FDR and recall is equivalent to sensitivity) through calculating the precision and recall for different thresholds of adjusted *P* values, and we empirically calculated the Area Under Precision-Recall Curve (AUPRC) value. We found ENIGMA (trace) performed consistently better than other methods according to the AUPRC metric (Figure S8c).

While the simulation study could give us a golden standard of CTS-DEGs that could help us to benchmark the algorithms, in realistic application, the CTS-DEGs inference would be influenced by many different factors (e.g. the number of latent cell types, the quality of reference expression profile). To show that ENIGMA has potential in identifying CTS-DEGs in realistic dataset, we used RNA-seq dataset of flow-sorted malignant cells, endothelial cells, immune cells, fibroblasts, and bulk samples from freshly resected human primary non-small-cell lung tumors to benchmark CTS-DEGs identification (n = 24 patients)[27]. To construct reference expression profile, we used a previously developed algorithm[8] to construct a batch effect corrected reference profile based on single cell NSCLC RNA-seq dataset from six donors[21] (Fig. 3e). Previous research and our principal component analysis (PCA) indicated that both malignant cells and fibroblast cells of lung squamous cell carcinoma (LUSC) and lung adenocarcinomas (LUAD) had distinct expression pattern differences[8, 27] (Figure S9). Therefore, we benchmarked our CTS-DEGs identification in these two cell types. We used concordance index based on previous work[8] to reflect the concordance between our predicted CTS-DEGs expression trend and ground truth and then generated its statistical distribution using a bootstrap method to evaluate the performance in identifying DEGs in a cell type-specific manner (Method). The result showed that expression concordance between ENIGMA-inferred CTS-DEGs’ CSE profile and flow-sorted expression profile was significantly higher than the concordance between bulk RNA-seq derived DEGs’ profile and flow-sorted expression profile (Fig. 3f, Figure S10a, b). Such results were further corroborated by scRNA-seq dataset of NSCLC tumors (Figure S10c, d). To make the analysis more comprehensive, we also defined the pseudo-ground truth CTS-DEGs based on flow-sorted gene expression profile and scRNA-seq with very strict criteria (|FoldChange| > 4 & FDR < 0.01). Based on this, we analyzed the CTS-DEGs identification sensitivity, and plotted its function as the number of top DEGs for each method. We found that both two versions of ENIGMA showed consistently higher sensitivity compared with all other algorithms (Fig. 3g). To further support the sensitivity of ENIGMA, we inspected the significances (*P* value) of those pseudo-ground truth DEGs of tumor cells in ENIGMA-inferred CSE and bulk RNA expression profile. The results showed that these genes were more significance in ENIGMA-sorted tumor expression profile compared with bulk RNA-seq dataset regardless up-regulated or down-regulated expression changes (Fig. 3h). These above results support that ENIGMA improves the identification of CTS-DEGs.

### ENIGMA reveals monocyte pseudo-differentiation trajectory in arthritis patients

We then explored the potential applications of ENIGMA for characterizing cellular heterogeneity in synovial biopsies from patients with arthritis. Rheumatoid arthritis (RA) is an autoimmune disease with chronic inflammation in the synovium of the joint tissue[28]. This inflammation leads to joint destruction, disability and shortened life span[28]. Studying the histological change from osteoarthritis (OA) to rheumatoid arthritis (RA), and defining key cell subpopulations with their activation states in the inflamed tissue are critical to define new therapeutic targets for RA[29]. Using flow-sorted RNA-seq profiles from RA to build a reference matrix[30], we inferred CSE at sample level to dissect four major cell types (includes Fibroblast, Monocyte, B cell, T cell) from the transcriptomes of 174 bulk RA & OA profiled by an independent study[31] (Fig. 4a). We benchmarked our *in-silico* purification through evaluating the CTS-DEGs expression change concordance with the external single cell RNA-seq datasets generated from synovial biopsies of patients with RA and OA[30]. In all four cell types, ENIGMA exhibited superior performance compared with other algorithms in relation to the DEGs expression concordance with the single cell RNA-seq dataset (Fig. 4b, Figure. S11a). Meanwhile, we found that using batch effect correction strategy could also improve the DEGs concordance, suggesting the importance to remove the confounder effect between reference and bulk expression matrix. Notably, we repeated our analysis by using reference matrix derived from single cell RNA-seq data from RA and observed similar results (Figure. S11b).

**Fig 4.**
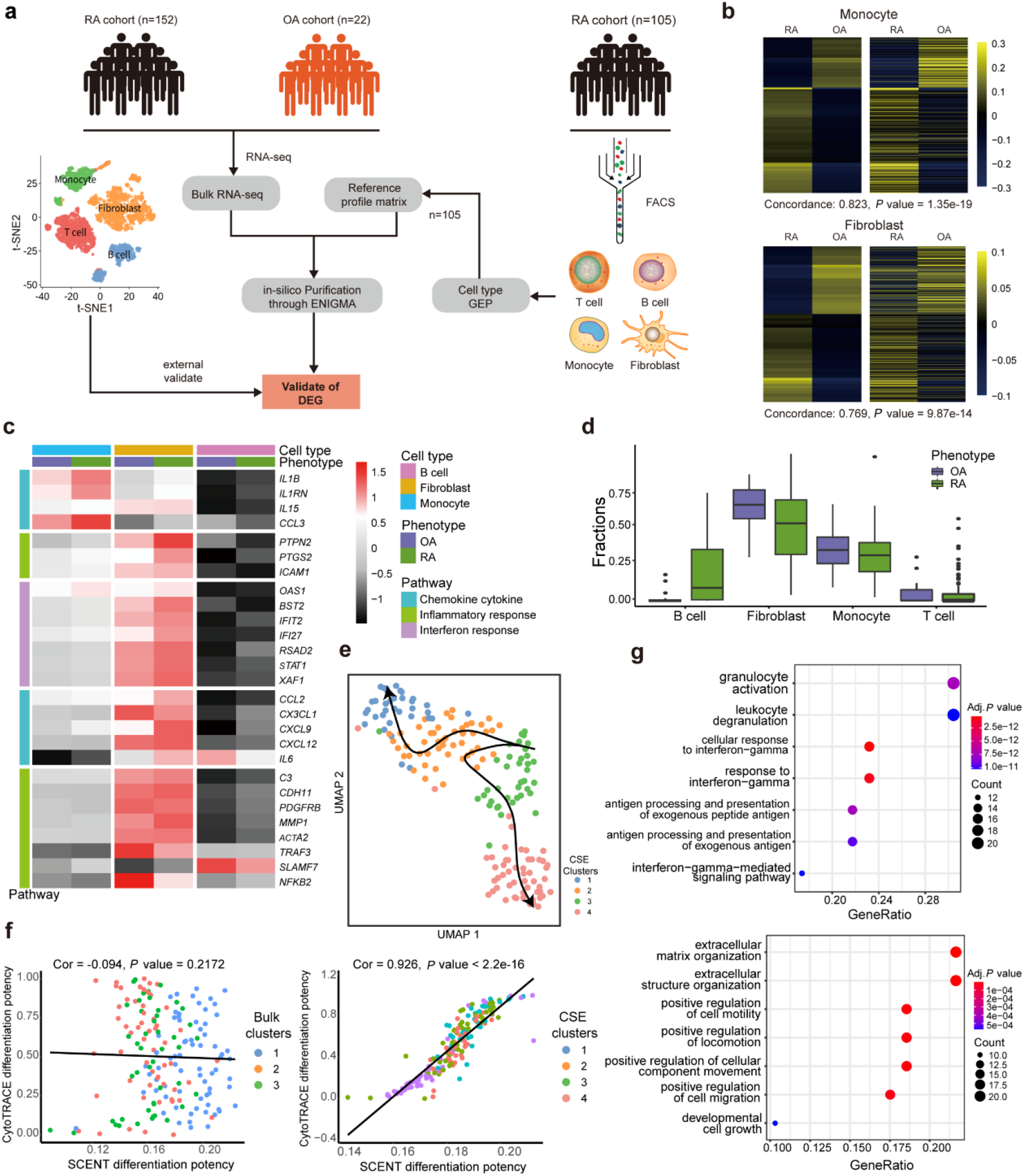
ENIGMA reveals monocyte pseudo-differentiation trajectory in arthritis patients. a. Schema outlining the application of ENIGMA to identify cellular signature in arthritis patient dataset. The t-SNE plot represents the external single cell RNA-seq data of arthritis patients for validation. We used one sample of flow-sorted RNA-seq (n = 105) to build signature (reference) matrix, and applied the signature matrix to deconvolute bulk RNA expression matrix (n = 174) by ENIGMA. To validate the reliability of *in-silico* purification by ENIGMA, we used scRNA-seq dataset and flow-sorted RNA-seq datasets (n = 104) to perform CTS-DEG external validation. b. Analysis of the concordance between ENIGMA-imputed (left) and scRNA-seq (right) CTS-DEGs expression profiles in OA and RA. The heatmaps show the averaged mean-centered log2-adjusted expression values for each DEG identified by ENIGMA. The DEGs were defined as the genes with |log2FC| > 2 and adjusted *P* value < 0.01. ‘DEG concordance’ denotes the fraction of genes with the same direction of differential expression between ENIGMA and scRNA-seq profile. The *P* value was calculated through performing Wilcoxon rank-sum test of concordance index (Method). c. The heatmap of averaged expression value of pathway marker genes from each group (OA & RA) across ENIGMA predicted CSE of B cell, Fibroblast and Monocyte. d. The cell type fraction boxplot of four cell types in arthritis patients. Each box was colored according to the phenotype information. e. A UMAP plot of normalized monocyte CSE profile imputed by ENIGMA for both OA and RA. The sample was clustered using Louvian algorithm on the KNN graph built based on monocyte CSE. Each dot represented a deconvoluted sample belongs to the monocyte and colored according to the cell clustering results. The principal curve was fitted through R package *Slingshot* [84]and the orientation of the curves are given by CytoTRACE and SCENT differentiation potency prediction. f. Scatter plots of CytoTRACE predicted differentiation potency (Y-axis) with SCENT predicted differentiation potency (X-axis) of observed bulk expression profile (left) and predicted monocyte CSE profiles (right). Each dot represented a sample and was colored according to the data clusters. g. The top seven significant GO pathways of the two branch-specific gene sets identified using the *tradeSeq [40]* through generalized additive model. Left and right panels showed the GO terms of the first (cluster 3 to cluster 4) and second (cluster 3 to cluster 1) branches, respectively. The pathways were ordered according to their statistical significance. The GO analysis was conducted using R package *clusterProfiler*.

Within *in-silico* purified cell subpopulations, we discovered many significant CTS-DEGs in monocytes and fibroblasts that could distinguish two disease states of arthritis (Fig. 4c). For instance, in fibroblasts, we found that a lot of interferon response associated genes were up-regulated in RA patients. In monocytes, we found that some chemokine and cytokine associated genes (e.g. *IL1B, IL1RN* and *CCL3*) were up-regulated in RA patients (Fig. 4c). They play a crucial role in homeostasis, generation of cellular and humoral immune responses, as well as pathologic immune contribution in rheumatoid arthritis[30, 32–34]. The abundance and activation of macrophages in the inflamed synovial membrane significantly correlates with the severity of RA[35]. In the RA synovial membrane, it is known that a pivot differentiation step is from immigrated monocytes to mature macrophages[36, 37]. Therefore, studying the sample-wise co-expression structure of purified CSE of monocytes may give us a deep understanding of pathogenetic stage of arthritis.

Using UMAP to visualize the CSE profiles of monocytes and applying Louvain clustering algorithm to cluster the CSE profiles, we observed a pseudo differentiation trajectory constituted by four clusters that may related to the pathogenetic progression of RA (Fig. 4e). We performed the same analysis in bulk expression data and found that raw bulk expression could not recover all of those four clusters (Figure S12). To determine the initial point of differentiation trajectory, we used two well-known differentiation potency scoring methods, CytoTRACE [38] and SCENT [39], to evaluate the differentiation state of each cluster. Strikingly, the predicted scores for CSE profiles of monocytes had high concordance (Spearman’s Correlation = 0.926, Fig. 4f, Figure S13a, b), and both algorithms indicated that the samples of cluster 3 had the highest differentiation potency. This high concordance still held even when we regressed out the variations of monocytes’ cell type fractions across samples (Spearman’s Correlation = 0.858, Fig. 4f, Figure S14). We also observed that the ribosome genes were highly expressed in cluster 3 (Figure S13a, b). Notably, we noticed that in bulk samples, if we applied SCENT and CytoTRACE to predict underlying sample differentiation score, the two predicted scores no longer showed high concordance (Spearman’s Correlation = −0.09, Fig. 4f). As bulk samples is the admixture of different cell types, this observation may implied that the expression-based predicted differentiation potency score would be biased by the cell type compositional change, and couldn’t represent any differentiation state of the underlying cell type. Using the predicted differentiation potency score, we determined the initial differentiation point and fitted the principal curve on the UMAP space, which led to the discovery that two pseudo-lineage branches existed in the monocyte CSE profile (Fig. 4e). To study the underlying biological process related to these two branches, we fitted Generalized Additive Model (GAM) [40] to identify the specifically expressed genes within those two pseudo-lineage branches, and performed GO enrichment test. Surprisingly, the first lineage (cluster 3 to cluster 4) highly expressed the genes related to leukocyte degranulation, interferon-gamma (IFN-γ) response and antigen processing/presentation (Fig. 4g). The second lineage (cluster 3 to cluster 1) highly expressed the genes related to the extracellular matrix organization, cell motility and migration, and developmental cell growth (Fig. 4g). It is known that during monocytes to macrophages differentiation, monocytes would be polarized towards two opposite states (the M1 or classical activation and the M2 or alternative activation states) by cues in their microenvironment. IFN-γ can differentiate macrophages into M1 state that promoting inflammation, secreting pro-inflammatory cytokines and chemokines, presenting antigens, participating in the positive immune response [41]. But M2 activation state would release anti-inflammatory cytokines to reduce inflammation, and produce extracellular matrix (ECM) components, angiogenic and chemotactic factors [42]. Previous researches also indicated that M2 macrophages would have increased random and chemotactic motility [43]. Those lines of evidence supported the notion that the two pseudo-lineages of monocytes were biologically meaningful. We also found several important genes which are functional relevant with monocyte to macrophage differentiation and M1/M2 state phenotype. For instance, the first pseudo-lineage (cluster 3 to cluster 4) highly expressed the tumor necrosis factor coding gene *TNF*, which encodes the main pro-inflammatory cytokines produced by M1 activation macrophage (Figure S13c). This pseudo-lineage also highly expressed *S100A8/S100A9* genes (Figure S15), which are important regulators for macrophage inflammation [44]. In the second pseudo-lineage (cluster 3 to cluster 1), we observed that some important monocyte-to-macrophage regulator coding genes were highly expressed (*CAV1* and *DAB2*) (Figure S15) [45, 46]. In addition, we observed that some M2 state related genes were also highly expressed in the second pseudo-lineage, including hemopoietic growth factor coding gene (*CSF1*) [47], glucocorticoids receptor coding gene (*NR3C1*) [48] and epidermal growth factor coding gene (*EGFR*) (Figure S13c) [49]. These lines of evidence suggested that ENIGMA could reveal the latent cell type-specific cell states, and their transitional relationship across samples.

### ENIGMA identifies a senescence-associated gene co-expression module in beta cells from T2D patients

ENIGMA could also help us to reveal cell type-specific co-expression modules, and measure their association with phenotypes in large patient cohort. To further demonstrate and evaluate ENIGMA, we extended application with a well-studied tissue, pancreatic islet, which plays important roles in metabolic hemostasis and whose dysfunction leads to diabetes[50]. We applied ENIGMA to bulk pancreatic islet RNA-seq samples from 89 donors to estimate proportion and CSE for each cell type[51]. Next, we characterized their associations with hemoglobin A1c (HbA1c) level, an important biomarker for type 2 diabetes (T2D)[52]. After constructing cell type reference matrix from published pancreatic islet scRNA-seq dataset from normal donors, we dissected bulk RNA-seq dataset into five CSE profiles at sample level (Fig. 5a).

**Fig 5.**
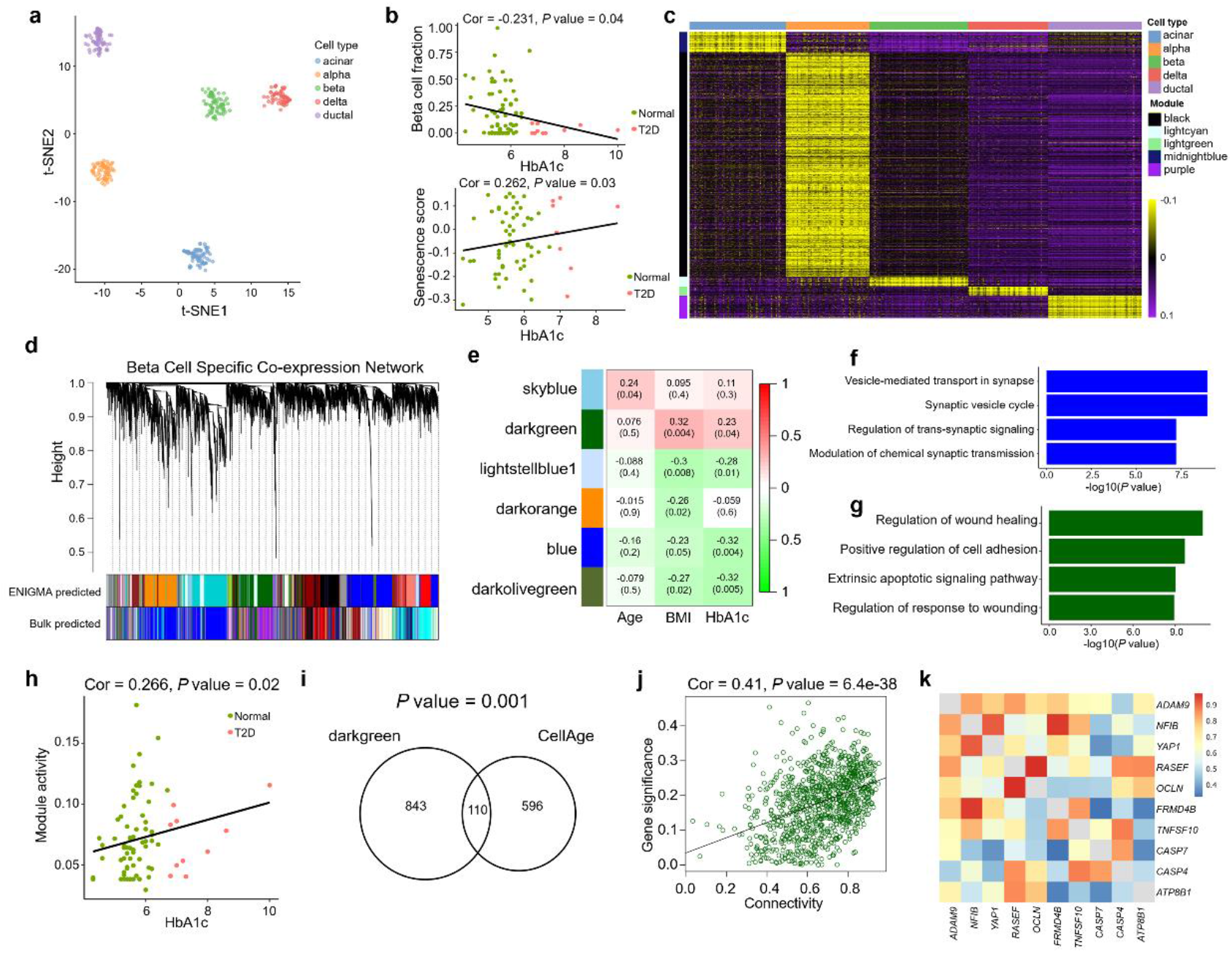
ENIGMA identifies cellular co-expression network in pancreas islet from T2D patients. a. A t-SNE plot of ENIGMA derived CSE of pancreatic islet bulk RNA profile sampled from T2D and normal patients. Each dot represented a deconvoluted sample and was colored according to the cell type it belongs to. b. The scatter plots of beta cell fractions (top) and senescence score (bottom) with the HbA1c level. The correlation was calculated using Pearson correlation coefficients. Each dot represented a deconvoluted sample. c. The heatmap of CSE of five co-expression modules that were associated with each cell type. The row represented a gene and the column represented a sample. d. Hierarchical cluster tree showing co-expression modules identified using WGCNA in beta cells. Modules were labelled by colors as indicated by the first color band underneath the tree. The modules identified by bulk RNA-seq were labelled by colors as indicated by the second color band underneath the tree. e. The heatmap of the association matrix between identified modules and three phenotypes (Age, BMI, HbA1c). The correlation coefficients were calculated using the Pearson correlation of each pair of the module eigengene expression value and phenotype. The module eigengene expression was calculated as the first left singular vector of the module gene expression matrix. f. The top four significant GO pathways enriched in blue module in panel e. The pathways were ordered according to their significance. GO analysis was conducted through R-package *clusterProfiler*. *g*. The top four significant GO pathways enriched in darkgreen module in panel e. The pathways were ordered according to their significance. GO analysis was conducted through R-package *clusterProfiler*. h. The scatter plot of darkgreen module (in panel e) activity with the HbA1c level. The correlation was calculated using Pearson correlation coefficient (Cor). Each dot represented a deconvoluted sample. i. Venn diagram showing the overlap of genes in darkgreen modules (in panel e) and genes in cell senescence signature database (CellAge)[57]. The significance of overlap was calculated using hypergeometric test. j. The scatter plot of gene significance (defined as the correlation coefficients between gene expression and HbA1c level, y-axis) and the gene connectivity (defined as correlation coefficients between gene expression and module eigengene expression, x-axis). Each dot represented a gene in darkgreen module in panel e. k. The heatmap visualization of topological overlap correlation network for top 10 hub genes from darkgreen module in panel e.

Pancreatic beta cells, which secrete insulin, are gradually lost during T2D[53, 54]. As our expectation, ENIGMA detected the significant negative association of beta cell proportion with HbA1c level (Fig. 5b, *P* value < 0.05). Recent works also suggested that insulin resistance induced the expression of aging markers, and beta cell aging (or senescence) could accelerate the progression toward diabetes[53, 55, 56]. We further asked whether ENIGMA could recover the latent senescence signal of beta cell in T2D patients. We constructed the expression-based senescence score depends on the senescence-associated genes downloaded from CellAge[57] and senescence-associated secretory phenotype (SASP) signatures[58] to predict the degree of senescence for each sample (Method), and applied the score on the beta cell CSE. We found that senescence score of beta cells showed a significantly positive correlation with HbA1c level and body mass index (BMI) (Fig. 5b, Figure S15, *P* value < 0.05), consistent with the previous finding that beta cell senescence increases with T2D and BMI[56]. To test whether senescence specifically happens in insulin secreting cell type, we applied senescence score on *in-silico* purified CSE and bulk RNA-seq samples (Figure S16). Notably, although we found that the senescence score in bulk RNA-seq samples showed positive correlation with HbA1c level and BMI (Figure S16), in CSE, only beta cells out of five cell types showed significantly positive correlation with both HbA1c level and BMI (Figure. S16). To show that this result could be reproduced, we constructed a second senescence score based on the senescence-associated genes downloaded from a different database (CSGene) [59] and the new senescence score of beta cells showed the strongest positive correlation with both HbA1c level and BMI than that of other cell types (Figure S17).

Gene co-expression networks capture biologically important patterns in gene expression data, enabling functional analyses of genes, discovery of biomarkers, and interpretation of phenotype associated biological variation. Previous network analyses were limited to the total gene expression levels in a whole tissue, while using ENIGMA could help us to deconvolute a mixing expression signal into a set of cell type-specific signals, further enable us to build cell type-specific gene co-expression network. We applied ENIGMA and weighted gene co-expression network analysis (WGCNA)[14] methods to find representative gene co-expression modules for each cell type. By performing network module discovery on all CSE, we identified several cell type-specific modules (Fig. 5c, Figure S18). For instance, in acinar cell, we identified the modules enriched with pathways like digestion and vitamin metabolic process, in line with the fact that pancreatic acinar cells are highly specialized exocrine factories that produce copious amounts of digestive enzymes for intestinal digestion[60]. We also identified synapse organization and synapse assembly pathways in delta cells, consistent with the idea that delta cells highly express the marker gene *GHSR* (ghrelin receptor), which is frequently involved in the process of synaptic organization and transmission [61]. These results support that ENIGMA could identify cell type-specific co-expression modules, making it possible to identify cell type-specific modules that related to the pathogenesis of complex diseases.

Next, we want to get into deep understanding about the dysregulation of beta cell specific pathways which are correlated with the T2D patients’ phenotypes. To make our analysis more reliable, we elaborated the single cell RNA-seq datasets of pancreas islet tissues from normal and T2D donors, to tune the model parameters. After inspecting the gene expression concordance between the inferred beta cell specific gene expression profiles and beta cell expression datasets from scRNA-seq, the parameter tuned ENIGMA model showed superior expression concordance with independent realistic dataset compared with other algorithms (Figure S19). Using the optimized deconvolution results, we built the gene co-expression network in beta cells to study their roles in the pathogenesis of T2D (Fig. 5d). We identified six modules in beta cells that correlated with one of the phenotypes (HbA1c, BMI or Age) and four of them (skyblue, darkgreen, blue and darkolivergreen) enriched with GO pathways (Fig. 5e). In blue and darkolivegreen modules, we found that vesicle-mediated transportation and cycling associated pathways were significantly enriched (Fig. 5f), and these two modules were negatively correlated with HbA1c and BMI (Fig. 5e), suggesting that both HbA1c and BMI factors may link to the deficiency of the secreting ability of beta cells. Meanwhile, we identified that the darkgreen module, which was positively associated with HbA1c and BMI, enriched a lot of pathways that associated with the regulation of apoptotic signaling pathways and wound healing (Fig. 5g), suggesting the dysfunction of beta cell in T2D patients. We also calculated the darkgreen module activity on all cell type-specific expression profile, and found that only in beta cell that this module was positively associated with HbA1c across all five cell types (Fig. 5h, Figure S20). We performed hypergeometric test and found that this module was significantly overlapped with senescence-associated gene signatures (Fig. 5i, *P* value = 0.001). Performing regression analysis between gene significance (which measured as the correlation significance with the HbA1c level) and gene connectivity (importance measurement of each gene in the module) showed a significant positive correlation, suggesting the hub genes of the module strongly connected to the HbA1c levels (Fig. 5j). The top 10 hub genes in the darkgreen module also recovered previously validated genes that play a role in T2D pathogenesis (Fig. 5k). For example, *TNFSF10*, which is part of TRAIL (tumor necrosis factor related apoptosis-inducing ligand), a type II transmembrane protein and a member of TNF-ligand family, has been extensively studied for its preferential ability to induce apoptosis of cancer cell[62, 63]. Growing lines of evidence support its involvement in the development of obesity and diabetes[64], and *TNFSF10* corporates with Caspase family protein to activate intrinsic mitochondrial pathway and cell apoptosis[64]. The hub genes also included some apoptosis and senescence-related cysteine peptidase (*CAPS7, CAPS3*)[65–67], further support the role of extrinsic signaling induced cell senescence and apoptosis in beta cell dysfunctions.

## Discussion

We developed a matrix completion-based algorithm, ENIGMA, for *in-silico* dissecting tissue bulk RNA-seq dataset through borrowing prior reference information derived from either FACS RNA-seq or scRNA-seq datasets. Comparing with previously proposed methods, ENIGMA used matrix completion strategy that jointly optimize all genes/samples’ reconstructed CSE profile, gave its capability to preserve or recover sample/gene correlation structure. Meanwhile, its great computational efficiency compared with previous algorithms make it possible to be applied on large cohort study. We conducted extensive benchmark on simulation datasets and application on realistic datasets to compare ENIGMA with state-of-the-art (SOTA) methods, demonstrating that ENIGMA improved the CSE estimation accuracy and recovered specific sample-wise correlation structure. Also, we validated that ENIGMA was relatively robust to the number of factors that may influence the inferred CSE quality compared with other SOTA methods. What’s more, the required running time of ENIGMA was dozens of times faster than other SOTA methods.

In addition, we showed that using ENIGMA could identify CTS-DEGs from bulk RNA expression profile, and demonstrated its advanced specificity and sensitivity compared with other algorithms. Nonetheless, ENIGMA also had limitations in CTS-DEGs prediction. For instance, for those cell types which had rare abundance in cohort, both maximum ℓ_2_ norm-based and trace norm-based ENIGMA would not conduct accurate estimation, thus influence the DEGs detection. Such issue could not be addressed by other methods either. Besides, ENIGMA was sensitive to the phenotype associated variations, therefore it tended to call more CTS-DEGs compared with other methods, which may suggest that it has relatively higher false discovery rate compared with some SOTA methods specifically designed for cell type-specific variation detection (e.g. CellDMC)[68]. But this drawback could be corrected by controlling the number of statistically significant DEGs (Figure S8).

Furthermore, ENIGMA enabled the identification of meaningful cell type-specific pseudo-trajectory that may help to explain the progression of complex diseases. In both our benchmark simulation and application on arthritis patients, we showed that ENIGMA could reconstruct the pseudo-trajectory of CSE (Figure S3e, Fig 4e). In contrast, we found that previously developed CSE inference methods could not capture the trajectory structure (Figure S3e). The power of preserving correlation structure also gave ENIGMA benefits to identify gene co-expression modules, and to perform association study with the phenotype data. With these features, we anticipated that ENIGMA would improve our understanding of the underlying mechanism of pathogenicity of complex diseases relying on specific cell types.

Same as ENIGMA, the cell type proportion of both bMIND and CIBERSORTx is fixed during the optimization procedure [8, 13], and the cell type proportion information is also estimated based on a reference-based model (NNLS or CIBERSORT). Reference-based deconvolution approaches rely on the availability of accurate references to estimate cell type proportions. The discrepancy (cross-platform) between the biology of the references and the bulk samples could introduce bias in estimated cell type proportions. To prevent the technique effect, we applied S-mode and B-mode methods to correct batch effect [8]. The applications on real datasets also supported that these correction methods could benefit the CSE inference (Figure S11), This question also could be the improvement direction for ENIGMA in the future, that is, how to implement an iterative procedure to realize estimation accuracy improvements on both CSE profiles and cell type fractions through maximizing the shared biological variation between reference profile matrix and each mixture sample. Another key factor that may influence the performance of ENIGMA was the choice of parameters. Fortunately, the short runtime of ENIGMA made the tuning of parameters more feasible. Users could tune their parameters through improving gene expression concordance between inferred CSE profiles and external single cell RNA-seq datasets (Figure S19). Besides, we also discussed the role of parameter *α* to inferred CSE profiles, providing useful information for parameter tuning (see Data processing and parameter analyses section of Supplementary Notes).

In summary, ENIGMA represented a broadly applicable algorithm that could be used to deconvolute bulk RNA-seq dataset into cell type-specific expression profile, helping researchers to study the underlying heterogeneity in tissues, and systemically determine correlation between CSE and a variety of phenotypes. Therefore, ENIGMA could be utilized as an efficient tool for biological discovery, and it provides a framework to perform integrative analysis on scRNA-seq and bulk RNA-seq dataset. We believe that this approach has great potential to enhance the analysis on multicellular organisms, and it will facilitate the applications of mixture RNA-seq in the era of cell type-specific expression study.

## Methods

### The mathematical model of ENIGMA

The input of ENIGMA includes (1) the bulk RNA-seq data 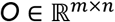, a count matrix represents expression profile from bulk RNA-seq data with columns representing *n* samples and rows representing *m* genes, and (2) the reference profile matrix, 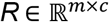, which represents the expression levels of *m* genes across *c* cell types of interest. Our goal is to completely deconvolute bulk expression data into series of cell type-specific expression matrix, (*X_i_*)_*i*=1, …, *c*_, in which 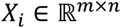, *c* represents the number of cell types.

In the previous study, Jiebiao Wang et al. proposed to use a full Bayesian model to reflect the relationship between bulk gene expression and cell type-specific gene expression[13].

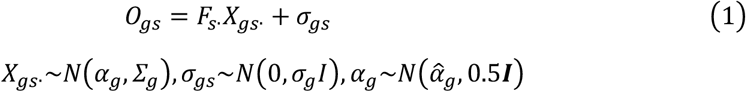

in which 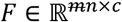 represents cell type fraction matrix, *X_gs_*. represents sample *s* level of gene *g* expression value across different cell types, and the parameter *∑_g_* represents the expression co-variance of gene *g* across different cell types. Jiebiao Wang et al. modeled each gene expression distribution across cell type *X_gs_*. as the Gaussian distribution, and the expected CSE expression profile *α_g_* ∈ *R^c^* was also modeled as the Gaussian distribution, and the expectation of the distribution was the average cell type-specific expression profile calculated from scRNA-seq, which they denoted as the profile matrix.

We borrowed the hypothesis raised by Jiebiao Wang et al. that the expectation of each CSE profile is the reference profile matrix, which can be learned from external scRNA-seq or FACS RNA-seq [13]. Besides, we assumed that the observation noise across different samples was the same. Taking those two assumptions into account, and inspired by previous imputation algorithm in scRNA-seq [69], we proposed to optimize the following loss function to estimate (*X_i_*)_*i*=1, …, *c*_

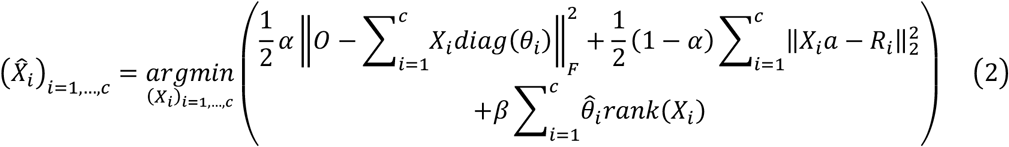

in which *θ_i_* represents 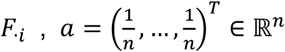, and *rank*(*X*) is the rank of the matrix *X*. The corresponding parameter *β* ∈ (0, +∞) controlling the latent dimensionality of each deconvoluted expression matrix. The rationality behind the rank norm regularization is that we think the rank of CSE matrix should be relatively smaller than bulk expression. Considering each CSE matrix represents the expression spectrum of one cell type, it would be more homogenous than bulk expression profile. Therefore, controlling the rank of CSE matrix would be a natural candidate regularization term as it measures the complexity or information content of the matrix. Meanwhile, different cell types would usually have different degrees of heterogeneity across tissues/samples which would be reflected in their CSE matrices. Therefore, constraining them with the same degree of penalty may lead to wrong estimation. We multiplied *β* with the expectation of *θ_i_*, *i.e*., 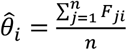, which means that if the average proportion of this cell type is low in bulk samples, the rank constraint will be relatively loose. In other words, there may have more differences for cell types with a small average proportion among individuals. The parameter 1 – *α*. (*α* ∈ (0,1)) is the weight for the agreement between the vector of aggregated CSE matrix and its corresponding reference expression vector (*R_i_*). Reasoning about why using *α* and 1 – *α* to weight object functions is described in the Loss design section of Supplementary Notes.

### Trace norm model

The rank regulation term constrains the complexity of the model, but unfortunately, except for a few cases, the optimization problems involving rank constraints are difficult to solve[70]. At this point, the trace norm model is a special example of the low-rank regularizer to solve this problem [71]. We employed this model to compute the optimal solution as follows:

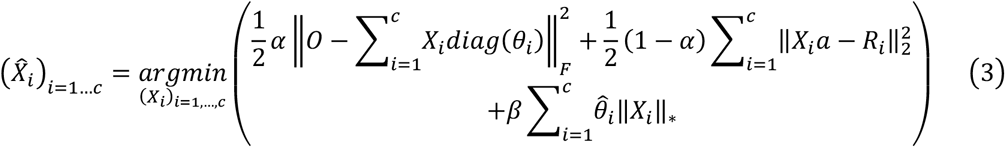

in which 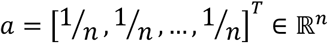. We used trace norm to regularize each decomposed expression profile, which is the convex envelope of rank function, calculated as the sum of the singular value of matrix. We used the following steps to calculate X in the above formula.

We transformed above constraint by introducing the auxiliary variable Y, and formulated optimization problem as augmented Lagrangian method as follow:

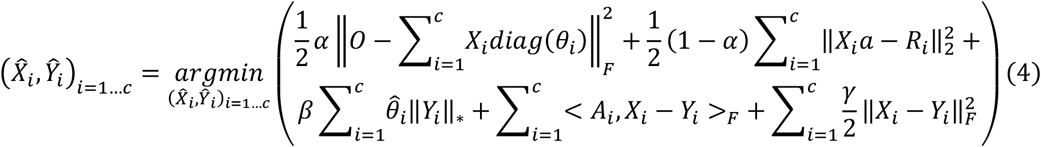

in which *γ*(*γ* > 0) is called the augmented Lagrangian parameter or step size.

The Alternating Direction Method of Multipliers (ADMM) [72] iteration scheme can be written as follows:

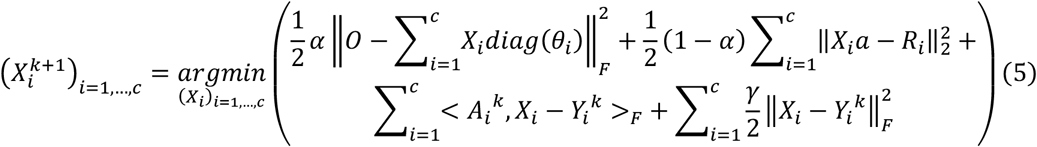

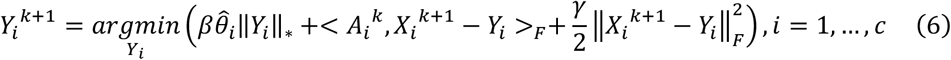

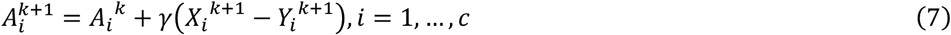

To simplify the equation, we use following new alphabet:

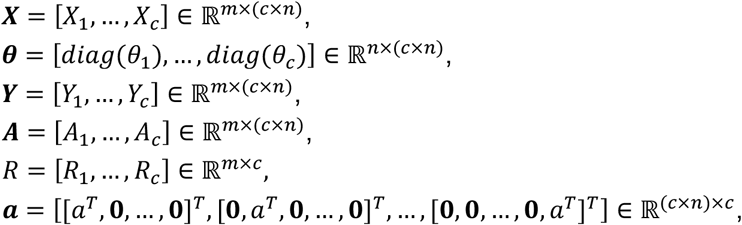

in which 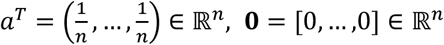

We could rewrite the first updated equation as follow:

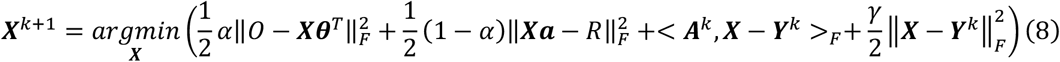

We took the derivative with respect to ***X*** and made it equal to 0 to get the equation as follow:

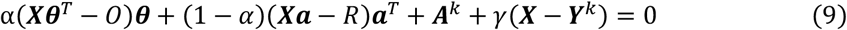

Therefore, the updated function of ***X*** could be written as follow:

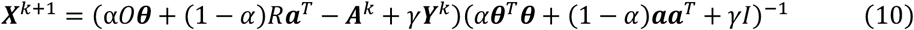

Further, we could separate the ***X***^*k*+1^ into *c* blocks to get the update of each CSE profile 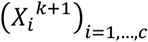 respectively, *i.e*., 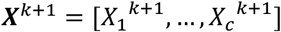.

For the optimization of (*Y_i_*)_*i*=1, …, *c*_, we could rewrite the equation as:

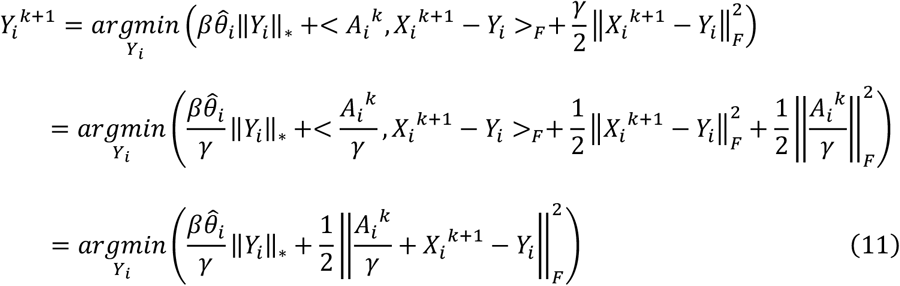

Further, we could use singular value thresholding [73] to calculate the 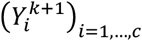

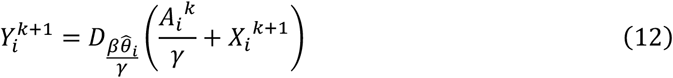

in which

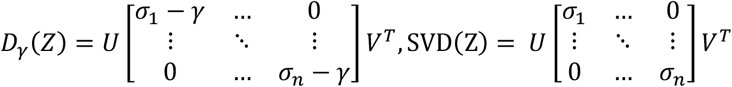

Finally, the updating of 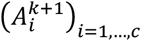 could be written as follow:

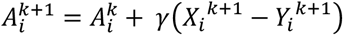

Putting all pieces together, we present an iterative way to address trace norm-based regularization problem (Algorithm 1).

#### Algorithm 1. D**E**co**n**volut**i**on based on Re**g**ularized **M**atrix Completion algorithm (trace norm version)

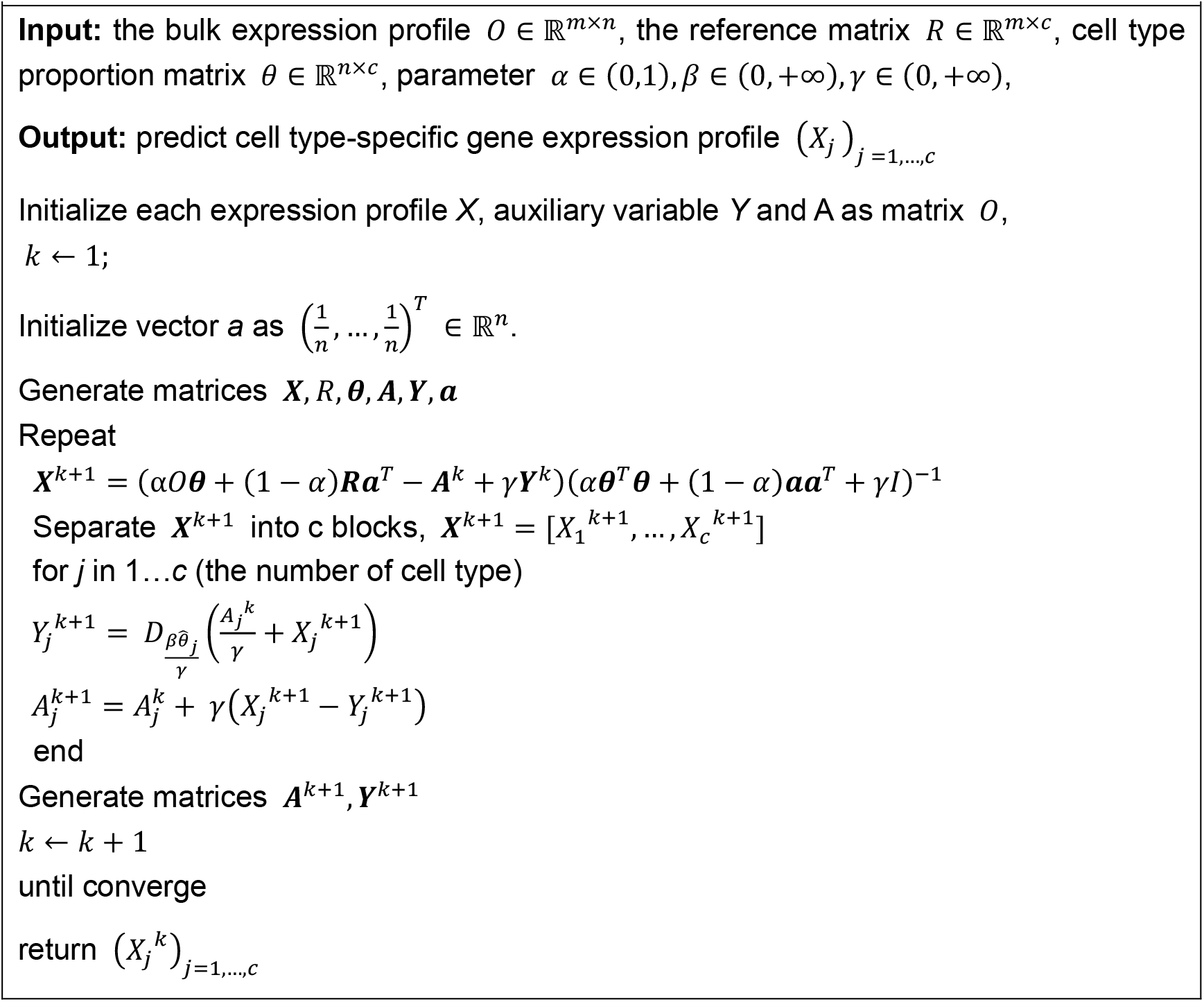

### Maximum ℓ_2_ norm model

The trace norm model has been successfully applied in many scenarios, but in our above derivations, it involves multiple singular value decomposition and matrix inversion, leading to high time complexity. At this point, it is easy to think of convex regularization term, the ℓ_2_ norm. For deconvoluting a bulk sample, we could use the following loss function:

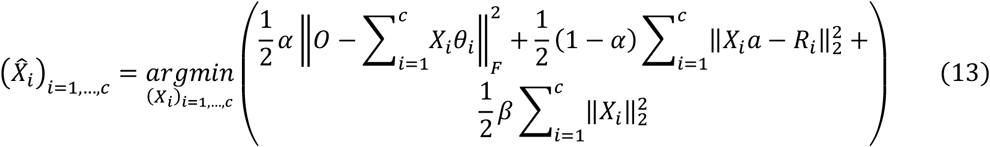

in which we regularized the deconvoluted expression vector through the ℓ_2_ norm, which penalizes the weight (deconvoluted expression) matrices from being too large. It was called ‘shrinkage’[74], which was frequently used in ridge regression or elastic net regression model to prevent overfitting. We could extend this formula to address multi-sample deconvolution as follow:

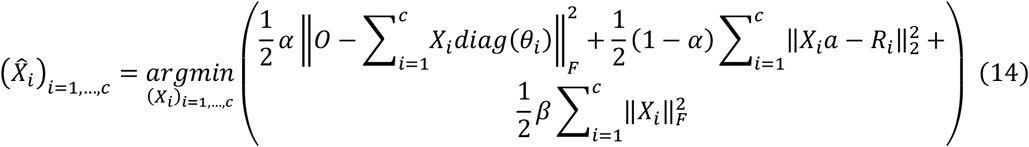

However, the F-norm regularization will constrain the overall expression value. We could release the constraint through considering the maximum ℓ_2_ norm across each column (sample) of deconvoluted cell type-specific expression matrices as follow:

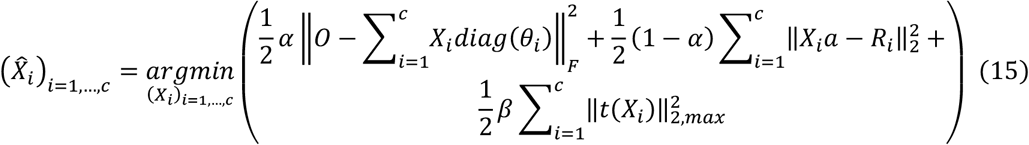

In which 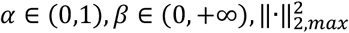 denotes the square of maximum ℓ_2_ row norm of a matrix:

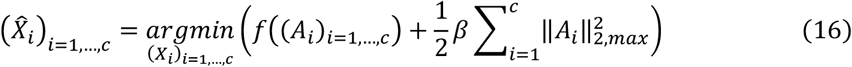

We calculated the partial derivative respect to each *A_i_*. Ideally, if we only consider the first term of the loss function, the iterate point could be represented as follow:

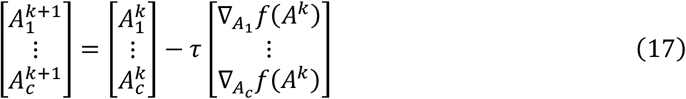

In which 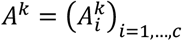. However, we required that the updated A needs to under the constraint of maximum ℓ_2_ norm. While it is hard to calculate this loss function gradient directly, we could approximate it. When performing gradient decent, we could use linearized proximal-point method [75] to replace the cost function *f* with a quadratic approximation localized at the previous iterate 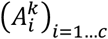.

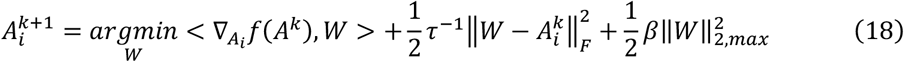

Where *τ* (*τ* > 0) is a proximal parameter, *W* ∈ *R^m×**n**^*. Then, we could complete the square to rewrite the formula:

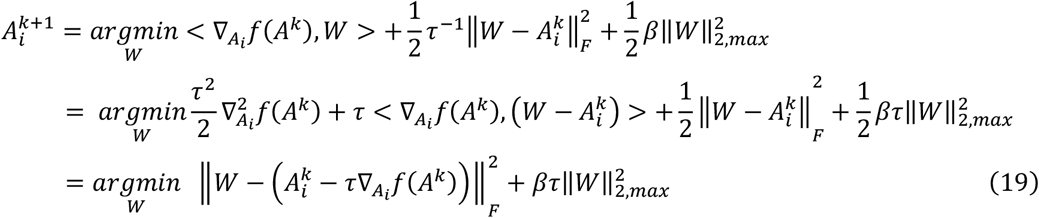

The new formula could be solved through previously proposed *squash* algorithm[75]. Which could be denoted as:

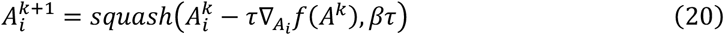

Putting all pieces together, we present an iterative way to address maximum ℓ_2_ norm-based regularization problem (Algorithm 2).

#### Algorithm 2. D**E**co**n**volut**i**on based on Re**g**ularized **M**atrix Completion algorithm (maximum ℓ_2_ norm version)

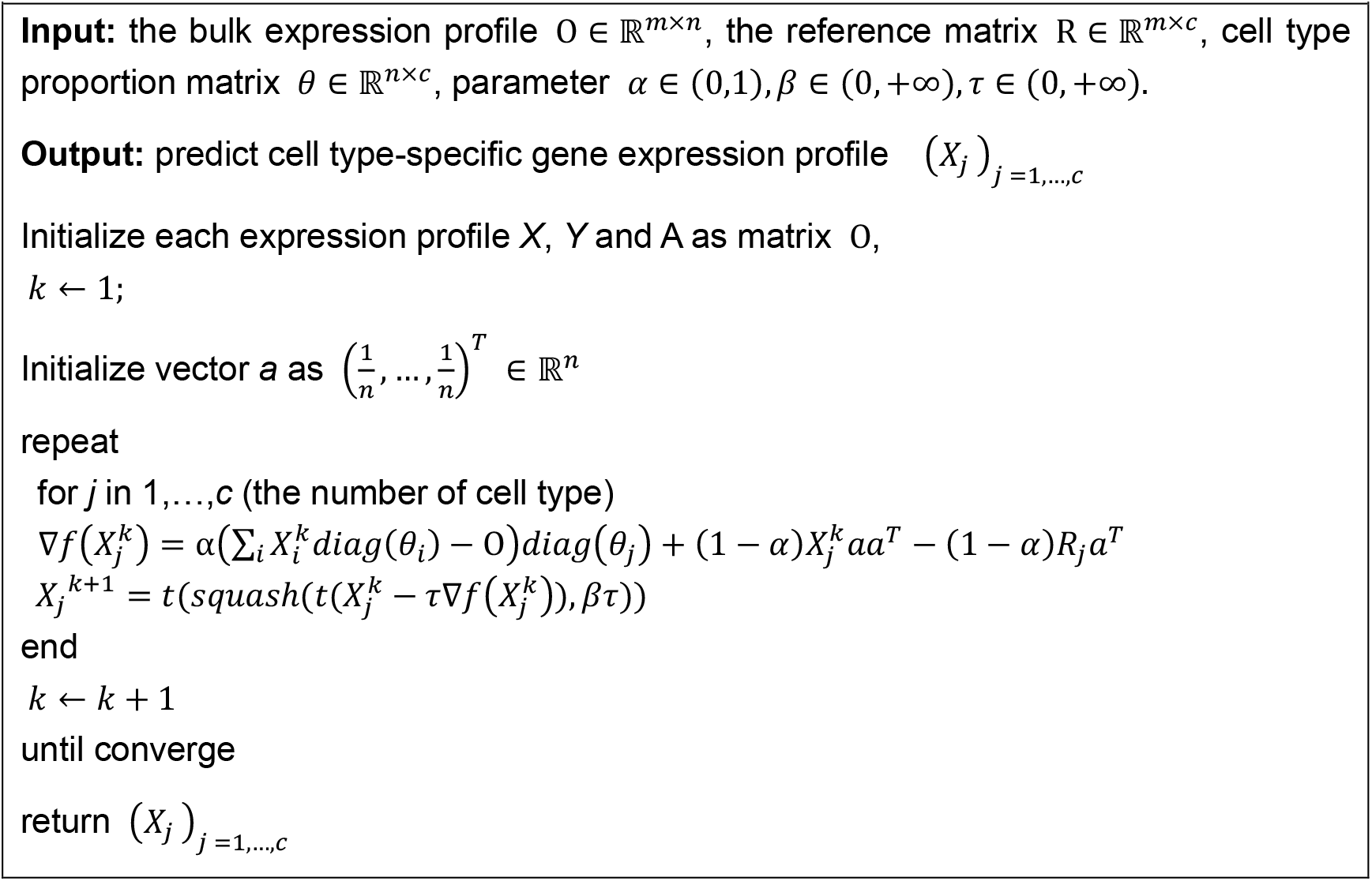

Different from trace norm-based regularization, this method would not promote rank reduced cell type-specific gene expression matrix, but the smoothed gene expression matrix with each entry of the gene expression matrix is shrunk. Also, training this model shall be faster than trace norm-based model, as its optimization procedure doesn’t involve the singular value decomposition or inversion of any matrix.

### Fundamental hypotheses of the two models

#### Less heterogenous hypothesis (trace norm model)

The inferred cell type-specific gene expression matrix represents the expression profile of a cell type, while the bulk expression represents the mixture of different cell types of heterogeneous tissues or samples, so there exists some variation in bulk expression driven by the latent cell type compositional change but not the gene expression alteration within a cell type. Therefore, it is natural to hypothesize that the CSE profile is less heterogeneous than bulk expression. We also used the bulk expression matrix as the initialization matrix of each cell type, which can ensure our inferred CSE matrices to have lower rank than bulk expression. Second, low-rank model is also widely used in gene expression imputation or prediction algorithms [69, 76], because it is universally known that in many biological processes, genes do not act in a solitary manner and rather interact with other genes to play the function. Those interactions make the expression levels of genes to be interdependent. The existence of gene collinearity can result in a highly correlated data matrix. So, assuming the gene expression values lie on a low-dimensional linear subspace, the resulting data matrix may be a low-rank matrix. Therefore, we used trace norm regularizer to promoting reduced rank cell type-specific gene expression matrix.

#### Hidden variable hypothesis (maximum ℓ_2_ norm model)

Most of cell type deconvolution algorithms, including ours, are reference-based deconvolution. Using reference-based methods could provide a robust and cost-effective *in-silico* way to understand the heterogeneity of bulk samples. It also assumes the existence of prior knowledge on the types of cells existing in a sample. These methods may fail to perform accurately when the data includes rare or otherwise unknown cell types with no references incorporated into the algorithm. Therefore, our reconstituted bulk expression profile (***Xθ**^T^*) may not include the variation from unknown rare cell types. Ideally, the observed matrix O would be more informative in our reconstituted bulk expression profile (***Xθ**^T^*). In other word, the observed bulk expression matrix would have higher rank than ***Xθ**^T^*. Under this hypothesis, we need to reduce the rank of ***Xθ**^T^*. We have proved mathematically that controlling the trace norm of reconstituted bulk expression profile (***Xθ**^T^*) equals to the maximum **ℓ**_2_ norm (***X***) of CSE profile (see loss design section of Supplementary Notes).

### Maximum ℓ_2_ norm model (ENIGMA) vs. trace norm model (ENIGMA (trace))

Both trace norm and maximum **ℓ**_2_ norm models show superior performance at different aspects. First, trace norm model poses trace norm regularizer to inferred CSE profiles, and uses low-rank matrix to approximate cell type-specific gene expression, which may help the model to discover better gene variation across samples. Trace norm could also perform better than maximum **ℓ**_2_ norm on CTS-DEGs identification (Fig. 3, Figure S8). Second, maximum ℓ_2_ norm has assumed that there exist unknown variables (expression of rare cell types or technique variations) in bulk samples, and maximum **ℓ**_2_ norm shows better performance on recovering cell type-specific correlation structure even there exists very strong noise in observed bulk expression matrix (Fig. 2d, Figure S3). So, choosing which model is dependent on what kind of analyses users want to conduct. When users want to define patients/samples subtypes according to cell type-specific gene expression profile (e.g. malignant cell), users could choose the maximum **ℓ**_2_ norm model to perform the deconvolution. Besides, when users want to perform cell type-specific analysis of differentially expressed genes, users could choose the trace norm model to perform the deconvolution. Maximum **ℓ**_2_ norm is also preferable if users have a large cohort of bulk samples. Finally, the training of maximum **ℓ**_2_ norm model is not involved with any inverse matrix calculation or singular value decomposition, so it is very scalable to the large bulk samples (Figure S6). When users want to perform fast deconvolution on the bulk expression dataset with large sample sizes, we suggest to use maximum **ℓ**_2_ norm model.

### Expression concordance estimation between ground truth expression profile and inferred CSE

Validate the DEGs prediction performance on real dataset is difficult because of the following considerations: 1) We don’t know the ground truth cell type-specific DEGs; 2) Different DEGs selection criterion (*P* value, FoldChange) would influence the performance evaluation. The key concern in expression deconvolution is that we want to know the gene expression variate or trend across case and control samples. Inspired by previous research[8], we calculated ‘DEG concordance’ as the fraction of genes within predicted DEGs with the same direction of differential expression between inferred CSE and the validation data (higher in both or lower in both) after mean-aggregating the expression data by known phenotypic groupings. To make the comparison of different methods more faithful, we randomly changed the predicted DEGs selection criterion as follow: First, we set a fixed threshold for *P* value or adjusted *P* value (< 0.05) to filter the genes, which was considered as tested significantly. Then we sampled the Fold Change (FC) cutoff on the FC distribution, and filtered the genes that > FC cutoff or < -FC cutoff, where these genes were considered as the predicted DEGs, and calculated their DEG concordance value. We repeated the selection of FC cutoff for 1000 times so that we could get a DEG concordance distribution. Using this method, we also could calculate the expression concordance on all genes, which measure the overall predicted trends similarity with the ground truth dataset.

### Construct reference cell type-specific gene expression profile

Building cell type-specific reference profile is necessary for cell type deconvolution. While the technical variation between bulk sample expression profiles and reference matrix will bring distortions in cell abundance and cell type-specific expression estimation, especially when we used scRNA-seq with technical dropout and noise. We applied previously developed methods, named as B-mode and S-mode normalization[8], to remove cross-platform batch effect.

B-mode normalization was used for the reference matrix derived from sorted bulk RNA-seq or microarray-based expression matrix. It also could be applied on the reference matrix derived from full length scRNA-seq with sufficient sequencing depth platform, e.g. SMART-Seq2[77]. S-mode normalization was used for the reference matrix derived from scRNA-seq platform with high technical dropout, e.g. 10X Genomics and droplet-based sequencing. S-mode normalization aimed to generate pseudo-bulk matrix through admixing the reference profile, and used ComBat[78] to correct the pseudo-bulk matrix with bulk expression matrix (which need to be deconvoluted), and restored the reference matrix through non-negative least square (NNLS) regression. We implemented both methods based on R and the source code could be found at https://github.com/WWXkenmo/ENIGMA.

### Running of other deconvolution algorithms

We also benchmarked cell type-specific expression profile inference methods TCA and bMIND packages. For TCA, we followed the tutorial provided by authors (https://cran.r-project.org/web/packages/TCA/vignettes/tca-vignette.html) and ran the algorithm with default parameter setting. One thing we need to note that this algorithm is originally designed for DNA methylation. In NSCLC real dataset analysis, we also used a simple analytic estimate solution of TCA and named it as “baseline” model. For bMIND, we followed the running tutorial https://github.com/randel/MIND/blob/master/MIND-manual.pdf. When running bMIND, we used the reference gene expression profile matrix of ENIGMA. All three methods need the cell fraction matrix (CFM), we therefore used robust linear regression with ad-hoc adjustment method[79] to estimate CFM.

We also compared ENIGMA with other methods when analyzing NSCLC datasets. These methods have intrinsic shortcomings thus are not suitable for performing extensive simulation study and benchmark. The first category is limited component method, ISOpure and DemixT, which only deconvolute the bulk RNA-seq dataset from tumor into two or three components (tumor cell or non-tumor cell), while the TCA, bMIND and ENIGMA could deconvolute multiple cell types. The second category is non-negative factorization method, CIBERSORTx, which is a web-server based tool, and time-consuming when applying it on large cohort. Also, it does not provide open source code for implementation. Thus, we used these three methods when performing NSCLC deconvolution, which is a bulk RNA-seq dataset sampling from NSCLC tumor with small sample size. We followed the benchmark workflow provided by CIBERSORTx[8], and applied Spearman correlation to measure the deconvoluted expression concordance with ground truth gene expression profile curated from corresponding FACS RNA-seq datasets.

### Data resource and analysis

#### Analysis of NSCLC

We used the Gentles et al. NSCLC bulk RNA-seq dataset [27] and Lambrechts et al. NSCLC scRNA-seq (10X Genomics) dataset [21] as the training datasets for predicting sample-wise cell type-specific expression profile. Then validate through NSCLC FACS cell type-specific RNA-seq [27]. For NSCLC scRNA-seq dataset, we first normalized the row count data as the count per million (CPM). This step was implemented through *Seurat* R-package [80] with R function NormalizeData(NSCLC_seurat, normalization.method = “RC”, scale.factor = 10000). Further, we constructed cell type reference expression profile based on NSCLC scRNA-seq dataset from Patient #3[21] through “S-mode” method described in the section “Construct reference cell type-specific gene expression profile” and set n_pseudo_bulk parameter as 1000. When estimating cell type fraction matrix, we used robust linear regression model based on our inferred reference expression profile and non-log transformed bulk expression profile. When performing bulk RNA-seq deconvolution to infer CSE, we used log transformed bulk expression and reference expression profile.

#### Analysis of rheumatoid arthritis

We used the Guo et al. synovial tissue bulk RNA-seq dataset sampled from rheumatoid arthritis and osteoarthritis patients as the discovery datasets [31]. We also used Zhang et al. FACS sorted RNA-seq dataset from four purified cell types (T cell, B cell, Monocyte and Fibroblast) as reference [30], and used its scRNA-seq (CEL-seq2) dataset as the validation dataset.

We processed the data as follow. To correct the batch effect while preserving biological variation between bulk RNA-seq dataset from whole synovial tissue and FACS RNA-seq dataset from four cell types, we used “B-mode” correction to construct the reference gene expression profile (as illustrated in ‘Construct reference cell type-specific gene expression profile’). We also transformed the scRNA-seq count matrix into CPM. To calculate differential expression genes (DEGs), we used Wilcoxon test to calculate DEGs across different cell types and used Benjamini–Hochberg method to correct multiple testing when calculating *P* value.

#### Analysis of pancreatic islet

We used Fadista et al. pancreatic islet sampled from type 2 diabetes and normal donors bulk RNA-seq dataset and corresponding patient metadata as the discovery dataset [51]. We also used Baron et al. pancreatic islet scRNA-seq dataset as the reference dataset [11], in which it only sampled the pancreatic islet from normal donors. Considering that the Baron et al. scRNA-seq dataset contains some rare cell types with no sufficient number of cells to generate pseudo bulk samples, we kept the cell type with its fraction in reference dataset more than 5%. We first normalized single cell gene expression data into CPM, and then used “S-mode” method to build cell type reference expression matrix. We used maximum **ℓ**_2_ norm model (ENIGMA) to perform deconvolution with *α* = 0.5, *β* = 0.1.

We performed weighted gene co-expression network analysis (WGCNA) [14] on purified CSE. We first analyzed CSE by concatenating all cell types. A signed weighted correlation network was constructed by creating a matrix of pairwise correlations between all pairs of genes across the predicted CSE. Further, the adjacency matrix was constructed by raising the co-expression measure, 0.5 + 0.5 × correlation matrix, to the power β = 10. The power value, which is interpreted as a soft-threshold of the correlation matrix, was selected through scale free fitness optimization routine [14]. Based on the resulting adjacency matrix, we calculated the topological overlap matrix (TOM), which is a robust and biologically meaningful measure of network interconnectedness [14]. Genes with highly similar co-expression relationships were grouped together by performing average linkage hierarchical clustering on the topological overlap. We used the Dynamic Hybrid Tree Cut algorithm to cut the hierarchal clustering tree, and defined modules as branches from the tree cutting. We summarized the expression profile of each module by representing it as the first principal component (referred to as module eigengene). Modules whose eigengenes were highly correlated (correlation coefficient above 0.85) were merged.

To tune the parameter of model for inferring beta cell specific expression profile more accurately, we calculated DEGs between T2D and normal controls on both inferred CSE and independent pancreatic islet single cell gene expression dataset [81]. We next compared the top 500 most significant differentially expressed genes expression concordance and all gene expression concordance of our prediction with that of independent scRNA-seq dataset. After parameter tuning, we used the CSE profiles inferred by optimized deconvolution model to build beta cell-specific weighted gene co-expression network. We first kept the top 8000 variable genes in beta cell CSE, then followed the above described way to construct the gene co-expression network step by step. We chose the power value 24 according to its scale free fitness result.

In senescence score assessment, we used two different ways to assess senescence. We curated two gene sets associated with cell senescence. One is SASP gene list [58] and the other is CellAge senescence-associated genes list [57]. To further support our results, we repeated the analysis based on a different gene set which is collected from CSGene database [59]. We then quantified expression variation of different gene sets across samples through GSVA (Gene set variation analysis)[82] and AUCell [83].

## Availability of data and materials

ENIGMA is implemented using R languages. The software packages are freely available under the MIT license. Source code has been deposited at the GitHub repository (https://github.com/WWXkenmo/ENIGMA) and Zenodo with the access code DOI: https://doi.org/10.5281/zenodo.2585902.

The analysis code and data used to analyze are available from the GitHub (https://github.com/WWXkenmo/ENIGMA_analysis) and Zenodo with the access code DOI: https://doi.org/10.5281/zenodo.2585885.

## Acknowledgements

This work was supported by National Key Research and Development Program of China [2018YFC1003500], National Natural Science Foundation of China [91949107, 31771336, 31521003] and Shanghai Municipal Science and Technology Major Project [2017SHZDZX01].

## Author contributions

The manuscript was written by W.W., X.Z., and T.N. and polished by T.N., Y.W., C.Z., W.T. and J.Z.. The method was conceived by W.W. X.Z. and J.Y. and the algorithm is implemented by W.W., X.Z. and J.Y. Computational analyses and algorithm evaluations were conducted by W.W. and X.Z. This work was supervised by T.N.

