## Supplementary Material for "Accurate estimation of cell-type resolution transcriptome in bulk tissue through matrix completion"

### Supplementary Materials

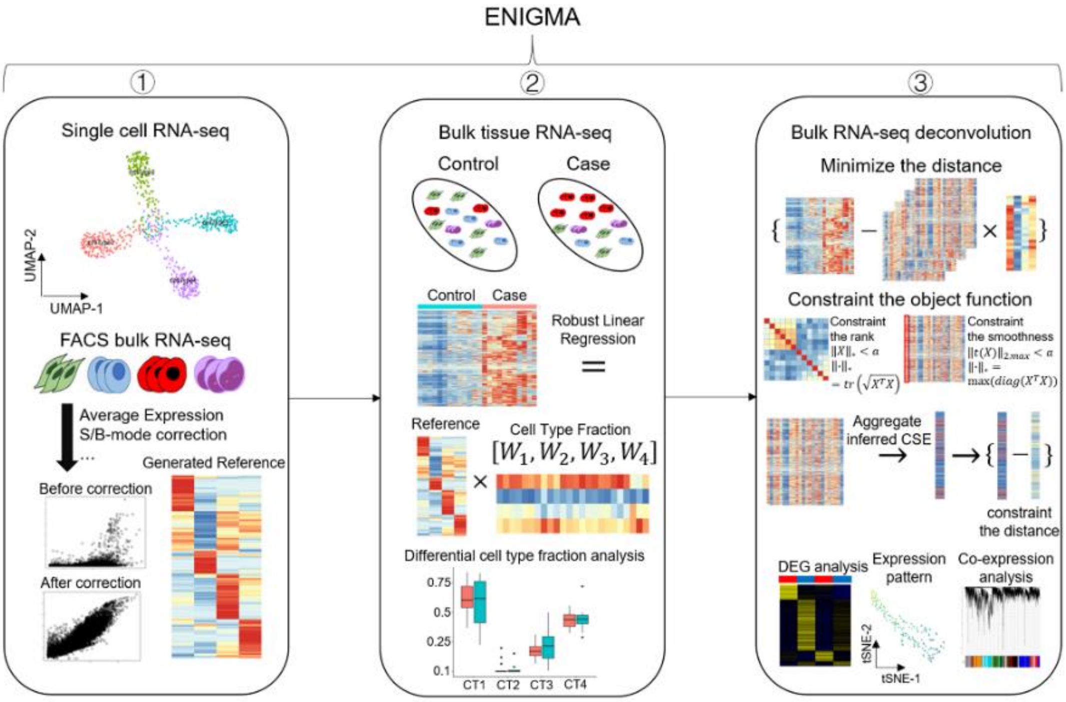


**Supplementary Figure S1. Workflow overview of the ENIGMA algorithm.**
The main steps of ENIGMA involve: 1) Transcriptome profiling of flow-sorted cell subpopulations or single cells to define a reference matrix consisting of features (genes) that can discriminate each cell subset of interest in a given tissue type. To prevent batch effect, ENIGMA applied pre-developed methods to correct the technique artifacts induced by cross-platform; 2) The bulk gene expression profile was deconvoluted through using robust linear regression and reference matrix to estimate each cell type fractions in each sample. The inferred cell fraction information could be used to perform differential cell fractions analysis; 3) Using matrix completion-based algorithm to deconvolute bulk RNA profile. There have two versions of ENIGMA, ENIGMA (maximum $\ell_2$ norm, default) and ENIGMA (trace norm). ENIGMA constrained that each average expression value inferred by CSE need to be close to the reference expression profile. After minimizing the object function to estimate the CSE, we could use the inferred CSE to perform differential gene expression (DEG) analysis, gene expression pattern analysis in each CSE, and gene co-expression analysis, etc.

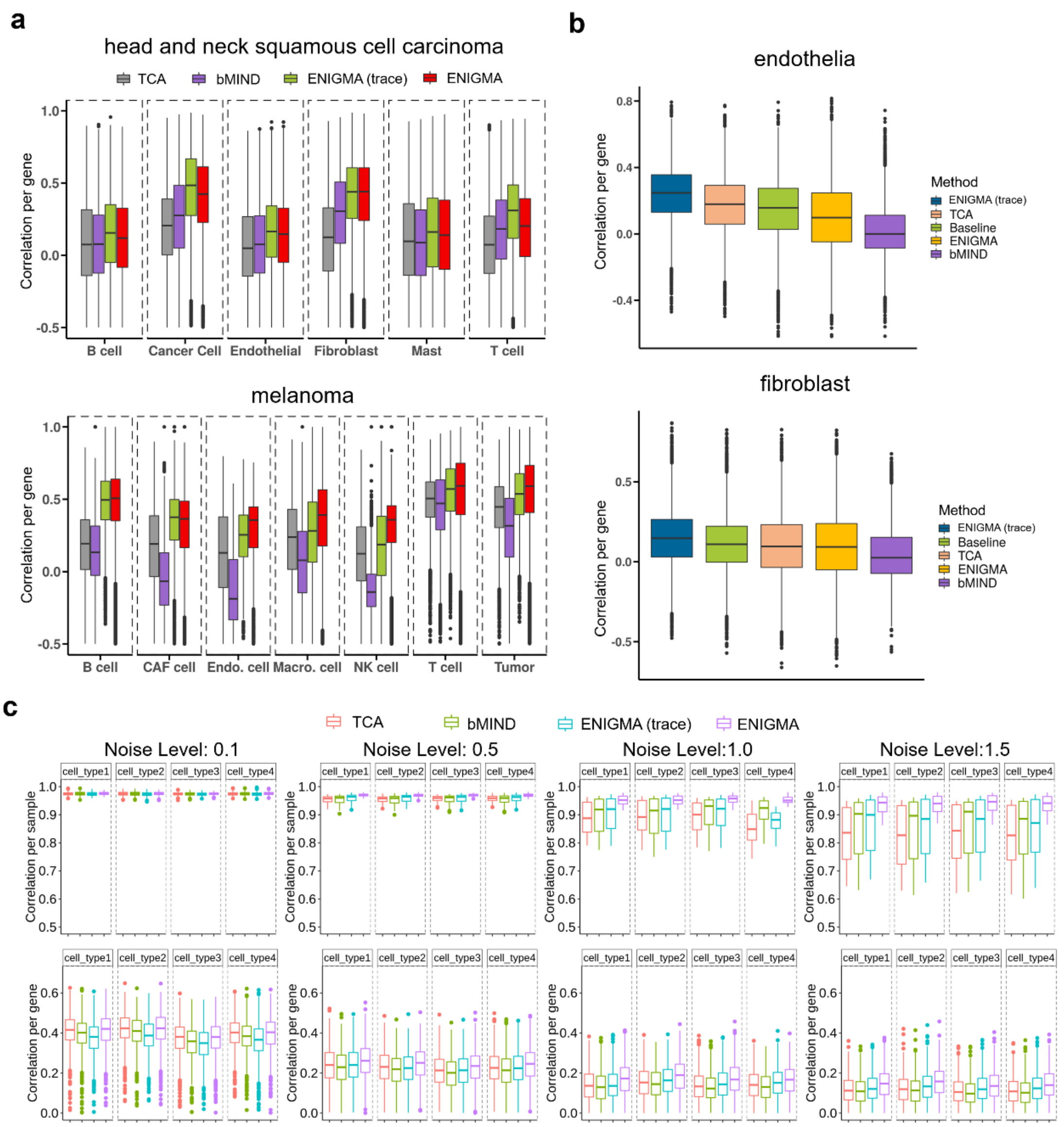


**Supplementary Figure S2. Benchmark of ENIGMA with other methods on realistic datasets.**

a. Each imputed CSE profile was quantile normalized and compared against its corresponding ground truth CSE by Spearman correlation. For each cell type, we computed correlation across samples for each gene (lower panel). The ground truth CSE and bulk expression profile for deconvolution was generated from scRNA-seq dataset of head and neck squamous cell carcinoma and melanoma.

b. The imputed NSCLC tumor specific profile was compared against its corresponding ground truth CSE by Spearman correlation. We computed correlation across samples for each gene (lower panel) in Endothelial (upper panel) and Fibroblast (lower panel). The ground truth CSE and bulk expression profile for deconvolution were curated from previous sequenced dataset of NSCLC [1]. The Spearman correlation was computed based on gene set filtered by CIBERSORTx. The *P* value was calculated from paired Wilcoxon rank sum test.

c. Each imputed cell type-specific expression (CSE) profile was compared against its corresponding simulated ground truth CSE by Spearman correlation with different observation noise level (0.1, 0.5, 1.0, 1.5). For each cell type, we computed correlation across genes for each sample (upper panel) and correlation across samples for each gene (lower panel). The CSEs were inferred through four methods, TCA, bMIND, ENIGMA, and ENIGMA (trace norm). The ground truth CSE and bulk expression profile for deconvolution were generated through simulation (Method).

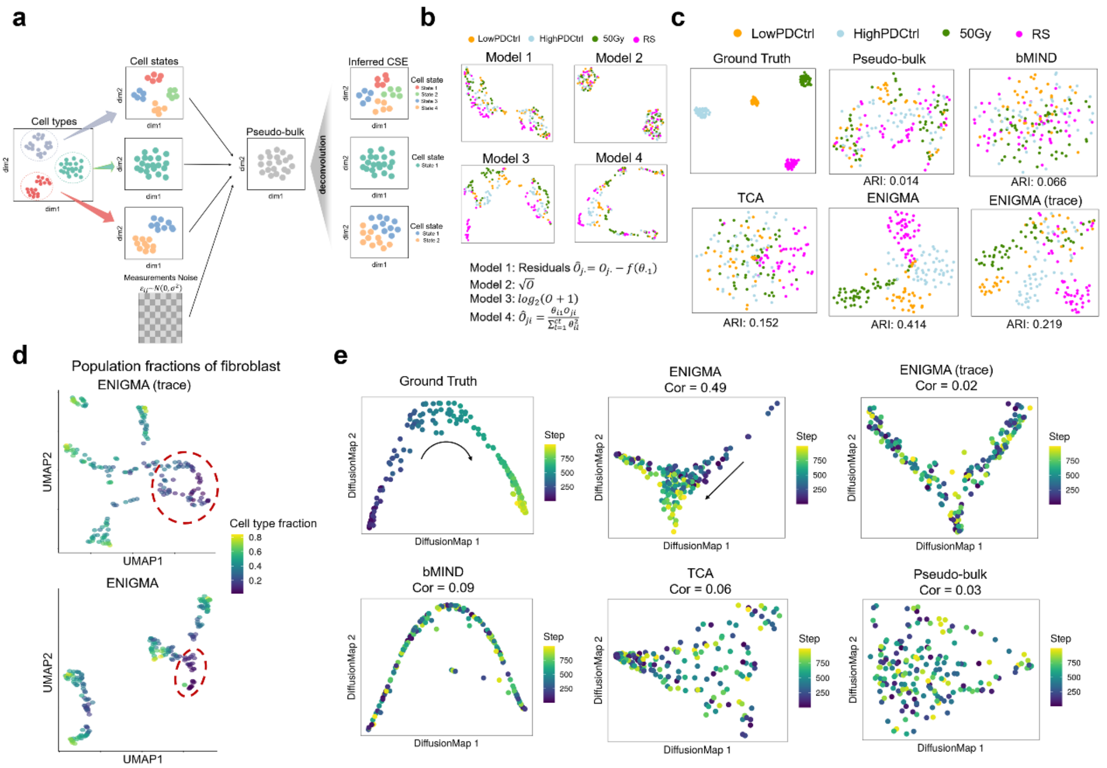


**Supplementary Figure S3. ENIGMA could recover the correlation structure among samples.**

a. Schema outlining the data generation model for benchmarking the performance of recovering latent cell states for each cell type. The bulk expression profile could be regarded as the linear combination of latent cell type-specific expression (CSE) profiles. Each CSE profile has specific clustering structure which encodes the transcriptome heterogeneity of this cell type across bulk samples. In our simulation model, for each of simulated cell type (three cell types in our simulation), we simulated its specific expression profile that containing latent clustering structure to group samples into distinct cell states. Then we aggregated CSE profiles and attached Gaussian white noise to create pseudo-bulk samples. A good CSE deconvolution algorithm is required to recovery the latent cell states information in its inferred CSE profiles. So, we clustered inferred CSE profiles deconvoluted from pseudo-bulk samples and used adjusted rand index to see if predicted cell states were consistent with ground truth cell states label. For the details about data simulation, please refer to Supplementary Notes. Benchmark tasks and metrics.

b. UMAP plots of simulated pseudo-bulk gene expression profiles processed by four RNA-seq processing methods. Each dot represented a bulk sample and colored according to the label of each bulk sample including fibroblast cell's senescence state. Model1: regress out the fibroblast cell fractions; Model2: sqrt transformation; Model3: log transformation; Model4: the baseline model proposed by Rahmani et al [2].

c. t-distributed stochastic neighbor embedding (t-SNE) plots of gene expression profiles. Each dot represented a simulated sample and was colored according to the fibroblasts senescence label (LowPDctrl (young quiescent cell), HighPDctrl (middle quiescent cell), 50Gy (X-ray induced senescent cells), RS (replicative senescent cell)).

d. Uniform Manifold Approximation and Projection (UMAP) plots of gene expression profile. Each dot represented a sample and was colored according to the fibroblast fractions.
e. Diffusion Map plots [3] of gene expression profiles. Each dot represented a sample and was colored according to the pseudo-time label on the right. The simulated ground truth CSE profile [4] was posed with a “trajectory” like structure (left of the upper panel), labeled as “Ground Truth”. The right of the bottom panel denoted ground truth profile mixed with other three simulated CSE profiles to generate a simulated bulk RNA expression profile, labeled as “Bulk”. We further used bMIND, TCA, ENIGMA and ENIGMA (trace) to deconvolute the admixture sample to infer the ground truth profile and inspect whether the algorithms recover the trajectory structure through visualization on DiffusionMap plot. To measure the performance, we also performed ordinary least regression of top two diffusion components with the pseudo-time label, and used the Pearson correlation between predicted pseudo-time value and ground truth pseudo-time value as the metric for benchmarking.

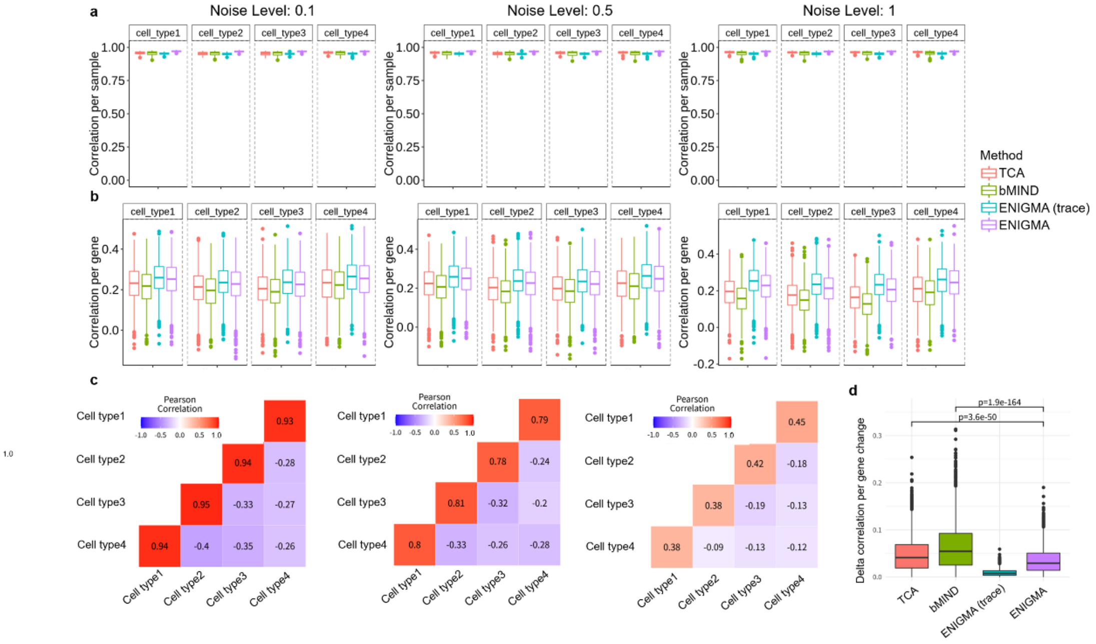


**Supplementary Figure S4. Benchmark of ENIGMA with other methods on simulated datasets with varying reference quality**.

a. Each imputed cell type-specific expression (CSE) profile was compared against its corresponding simulated ground truth CSE by Spearman correlation with different noise levels of reference profile (0.1, 0.5, 1). For each cell type, we computed correlation across genes for each sample. The CSEs were inferred through four methods, TCA, bMIND, ENIGMA, and ENIGMA (trace norm). The ground truth CSE and bulk expression profile for deconvolution were generated through simulation (Method).
b. Each imputed cell type-specific expression (CSE) profile was compared against its corresponding simulated ground truth CSE by Spearman correlation with different noise levels of reference profile (0.1, 0.5, 1). For each cell type, we computed correlation across samples for each gene. The CSEs were inferred through four methods, TCA, bMIND, ENIGMA, and ENIGMA (trace norm). The ground truth CSE and bulk expression profile for deconvolution were generated through simulation (Method).
c. The correlation between inferred cell type proportions and ground truth cell type proportions.
d. The boxplot of magnitude of correlation coefficient decrease. The *P* value was calculated through Wilcoxon rank-sum test. Less decreasing value meant better robustness.

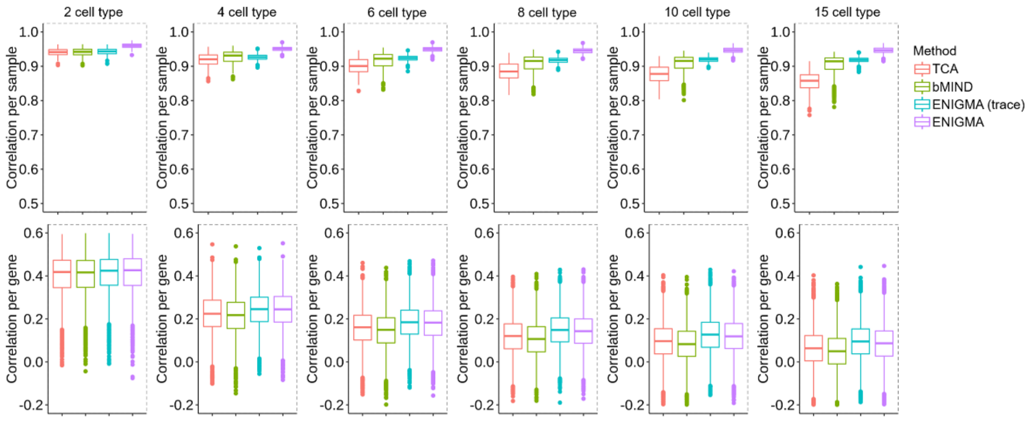


**Supplementary Figure S5. Benchmark of ENIGMA with other methods on simulated datasets with different number of latent cell types.**

Each imputed cell type-specific expression (CSE) profile was compared against its corresponding simulated ground truth CSE by Spearman correlation with different number of latent cell types (2, 4, 6, 8, 10, 15). For each cell type, we computed correlation across genes for each sample. The CSEs were inferred through four methods, TCA, bMIND, ENIGMA, and ENIGMA (trace norm). The ground truth CSE and bulk expression profile for deconvolution were generated through simulation (Method).

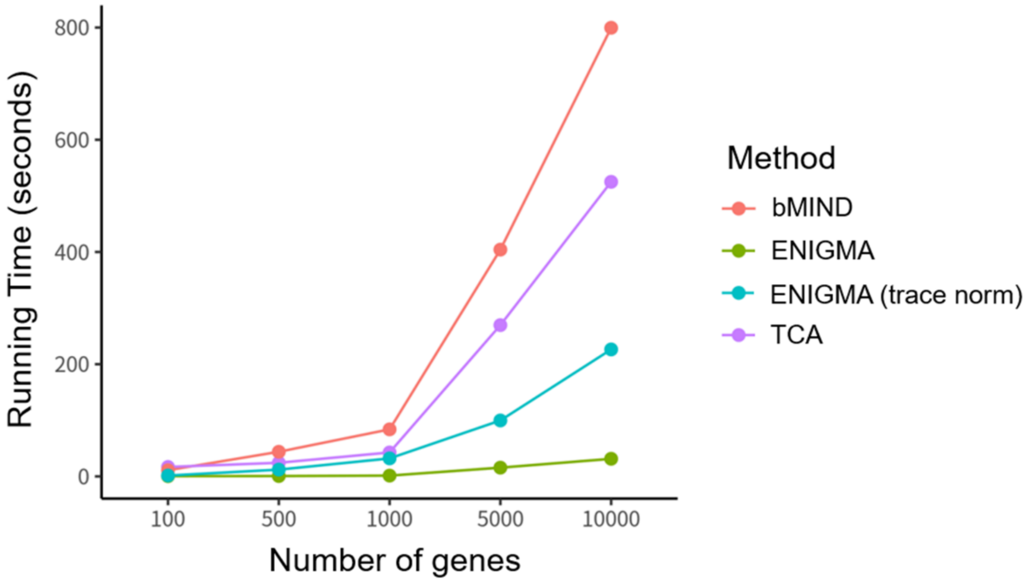


**Supplementary Figure S6. Runtime comparison among four methods.**
The scatter plot of running time variation according to the number of genes that need to be deconvoluted. Each dot and line were colored according to the method.

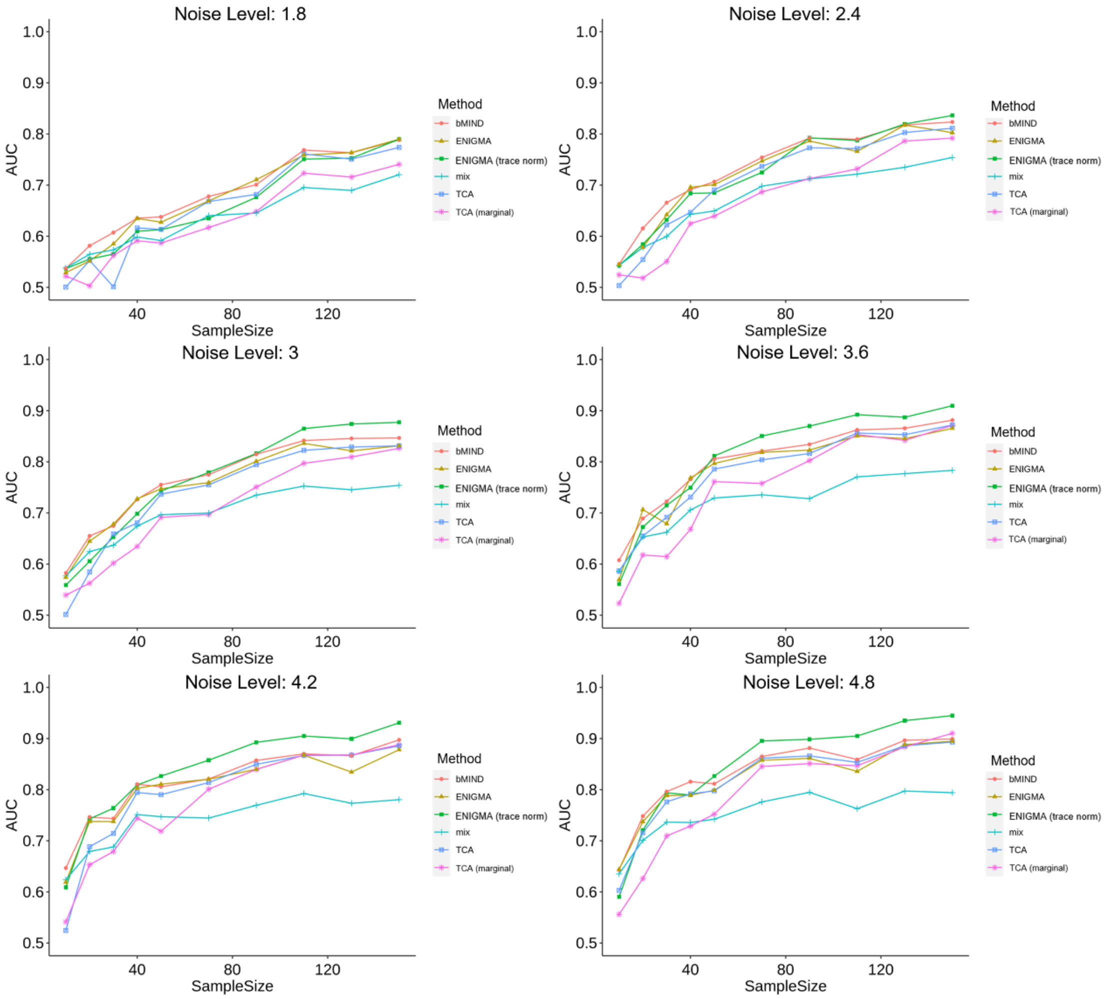


**Supplementary Figure S7. Area under the receiver operating characteristic curve (AUROC)-based power analysis with varying noise levels and sample sizes.** Scatter plots showing the mean area under the curve (AUC) for the recovery of known cell type-specific DEGs in 5 cell types with 10 runs of simulation as a function of the sample size of simulated bulk expression profile. We repeated the analysis with different signal to noise levels (1.8, 2.4, 3, 3.6, 4.2, 4.8).

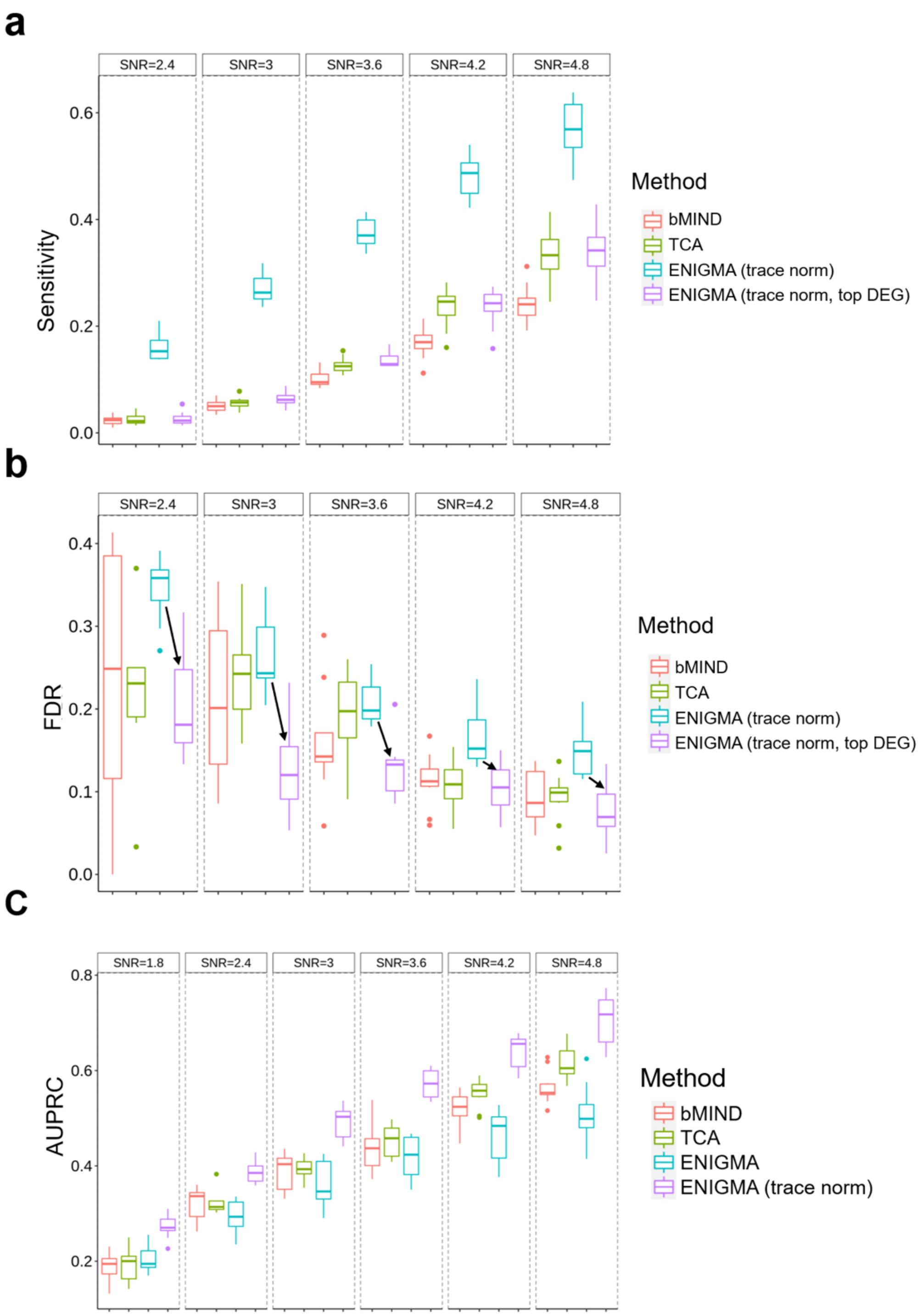


**Supplementary Figure S8. Benchmark inferred CSE on CTS-DEGs detection using FDR controlled DEGs calling method.**

a. Boxplots of sensitivity for the evaluation of detecting cell type-specific differentially expressed genes (CTS-DEGs) on the simulation dataset. The rate of true positives (sensitivity) was measured under a range of signal to noise ratio (SNR). All of the simulations were evaluated under the setting of five constituting cell types, 1000 genes, 100 samples and Gaussian observation noise with unit variance. Each cell type had 100 DEGs with randomly down- or up-regulated.
b. Boxplots of sensitivity for the evaluation of detecting CTS-DEGs on the simulation dataset. The false discovery rate was measured under a range of signal to noise

ratio (SNR). Y-axis denoted the FDR rate of top DEGs (the top *K* DEG genes, *K* equals to the number of DEGs identified by TCA).

c. Boxplots of AUPRC for the evaluation of detecting CTS-DEGs on the simulation dataset. The area of precision-recall curve was measured under a range of signal to noise ratio (SNR).

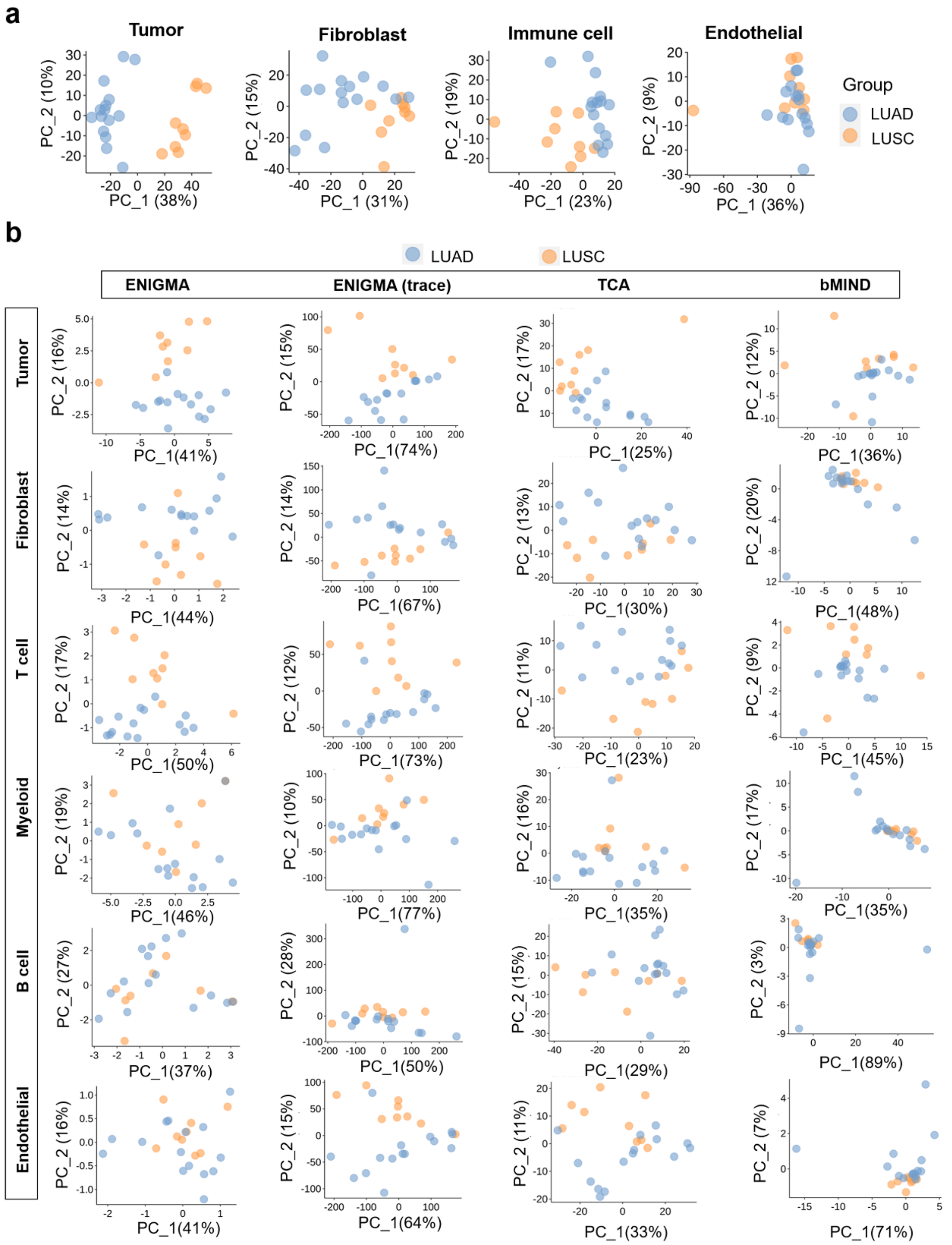


**Supplementary Figure S9. Validation of histology-specific DEGs in NSCLC tumors.**

a. The principal component analysis (PCA) plots of gene expression profile. Each dot represented a sample and was colored according to the tumor subtype (LUSC(1) or LUAD(0)).

b. The principal component analysis (PCA) plots of imputed CSE produced by four deconvolution methods. Each dot represented a sample and was colored according to the tumor subtype (LUSC(1) or LUAD(0)).

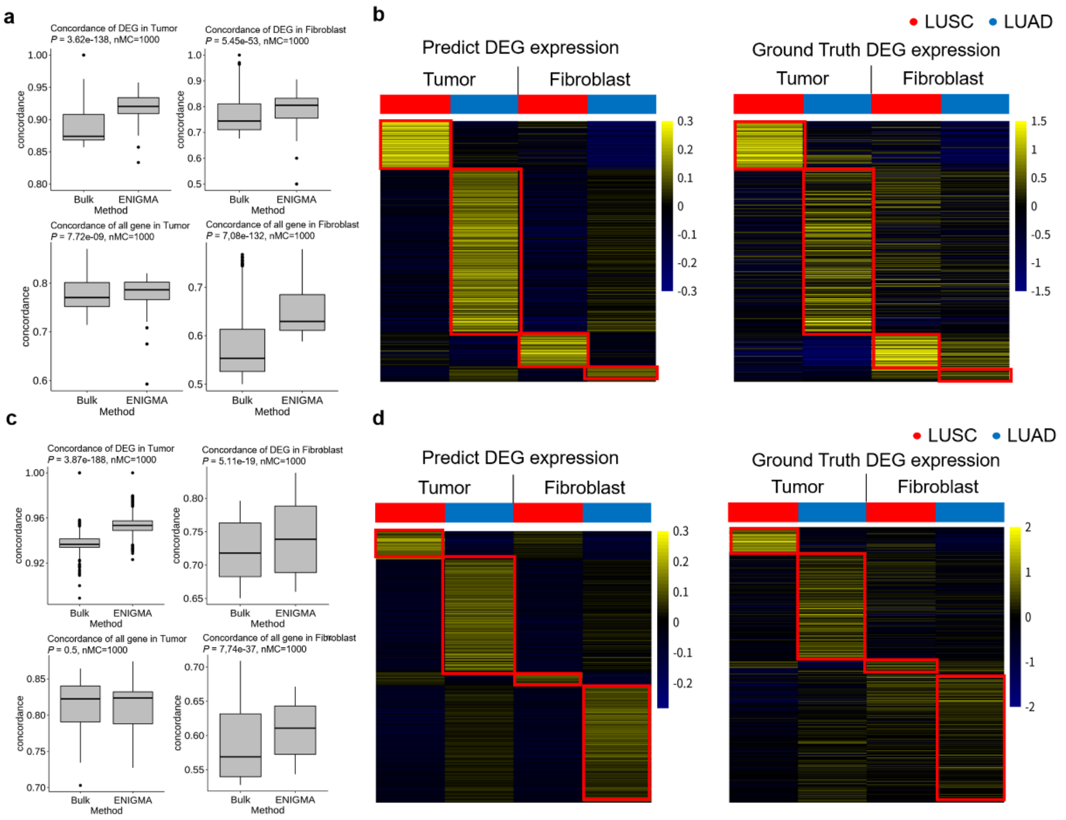


**Supplementary Figure S10. Validation of histology-specific DEGs in NSCLC tumors.**

a. The boxplots of gene differential expression concordance between CSE profile inferred from ENIGMA and flow-sorted RNA expression profile. The phenotype was the histological type of different NSCLC patients (LUSC vs LUAD). To make the comparison, the differential expression gene concordance between bulk RNA expression and flow-sorted RNA expression was also calculated. The gene differential expression concordance was calculated on all genes (upper panel) or DEG genes (adjusted *P* value <0.05, lower panel). The DEGs were defined on each expression profile respectively (bulk RNA expression profile or ENIGMA estimated cell type-specific expression profile). The *P* value was calculated through Wilcoxon rank-sum test. The reference matrix was constructed from a sample of flow-sorted RNA expression profile which was excluded in expression concordance calculation.

b. Heatmaps showing DEGs identified in epithelial/cancer, fibroblast populations identified in NSCLC tumors from CSE profile (left, n = 26 tumor samples) and RNA-seq profiles of corresponding FACS-purified populations from 26 patients with NSCLC of subtype LUAD or LUSC (right). The same genes were shown in both heatmaps and were ordered identically. For clarity, we visualized those DEG genes which were overexpressed in the specific cell type. Median centering was applied to each cell population separately. The reference matrix was constructed from a sample of flow-sorted RNA expression profile which was excluded in expression concordance calculation.

c. The boxplot of gene differential expression concordance between CSE profile inferred from ENIGMA and flow-sorted RNA expression profile. The phenotype was the histological type of different patients (LUSC vs LUAD). To make the comparison, the gene differential expression concordance between bulk RNA expression and flow-sorted RNA expression was also calculated. The gene differential expression concordance was calculated on all genes (upper panel) or DEGs (adjusted *P* value <0.05, lower panel). The DEGs were defined on each expression profile respectively (bulk RNA expression profile or ENIGMA estimated cell type-specific expression profile). The *P* value was calculated through Wilcoxon rank-sum test. The reference matrix was constructed from NSCLC single cell RNA expression profile which was excluded in expression concordance calculation.
d. Heatmaps showing DEGs identified in epithelial/cancer, fibroblast populations identified in NSCLC tumors from CSE profile (left, n = 26 tumor samples) and RNA-seq profiles of corresponding FACS-purified populations from 26 patients with NSCLC of subtype LUAD or LUSC (right). The same genes were shown in both heatmaps and were ordered identically. For clarity, we visualized those DEGs which were overexpressed in the specific cell type. Median centering was applied to each cell population separately. The reference matrix was constructed from NSCLC single cell RNA expression profile which was excluded in expression concordance calculation.

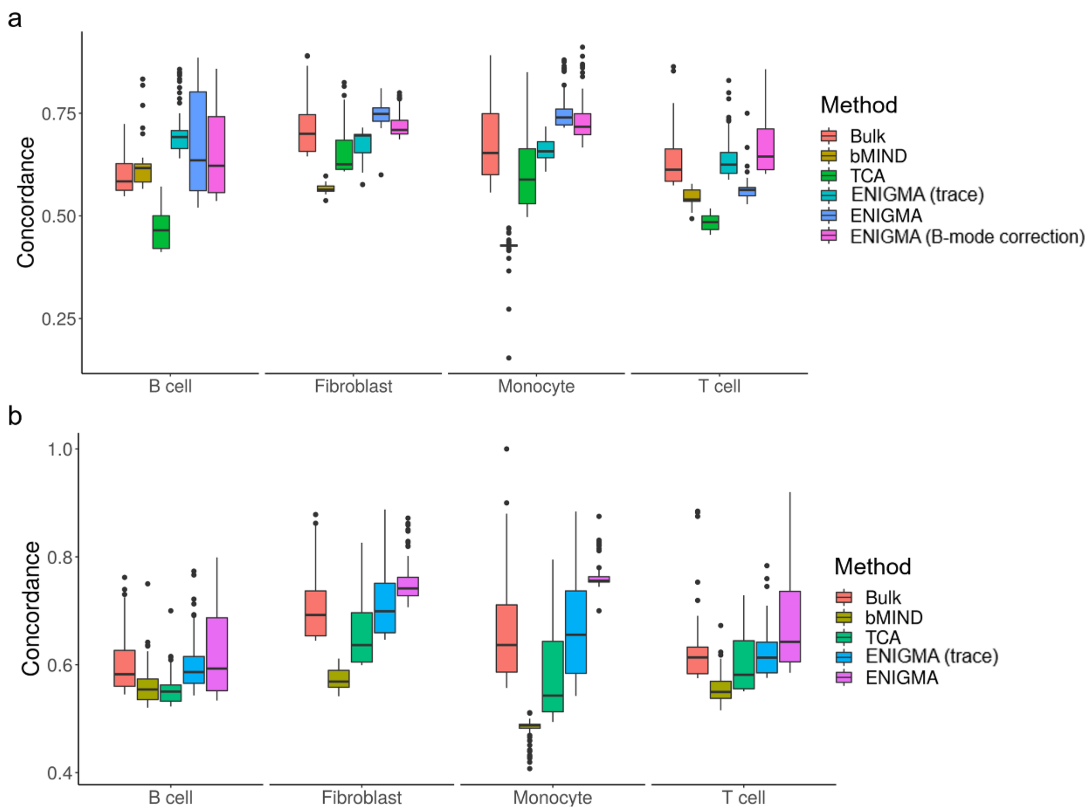


**Supplementary Figure S11. Validation of histology-specific DEGs in arthritis patients.**

a. The boxplots of gene differential expression concordance between CSE profile inferred from TCA, bMIND, ENIGMA, ENIGMA (B-mode correction), ENIGMA (trace norm) and single cell RNA expression profiles. The phenotype was the histological type of different arthritis patients (OA vs RA). To make the comparison, the gene differential expression concordance between bulk RNA expression and single cell RNA expression was also calculated. The gene differential expression concordance was calculated on DEGs (adjusted *P* value <0.05). The *P* value was calculated through Wilcoxon rank-sum test. The reference matrix was constructed from flow-sorted RNA expression profile.
b. The boxplots of gene differential expression concordance between CSE profile inferred from TCA, bMIND, ENIGMA, ENIGMA (trace norm) and single cell RNA expression profiles. The phenotype was the histological type of different arthritis patients (OA vs RA). To make the comparison, the gene differential expression concordance between bulk RNA expression and single cell RNA expression was also calculated. The gene differential expression concordance was calculated on DEGs (adjusted *P* value <0.05. The *P* value was calculated through Wilcoxon rank-sum test. The reference matrix was constructed from rheumatoid arthritis single cell RNA expression profile through S-mode correction.

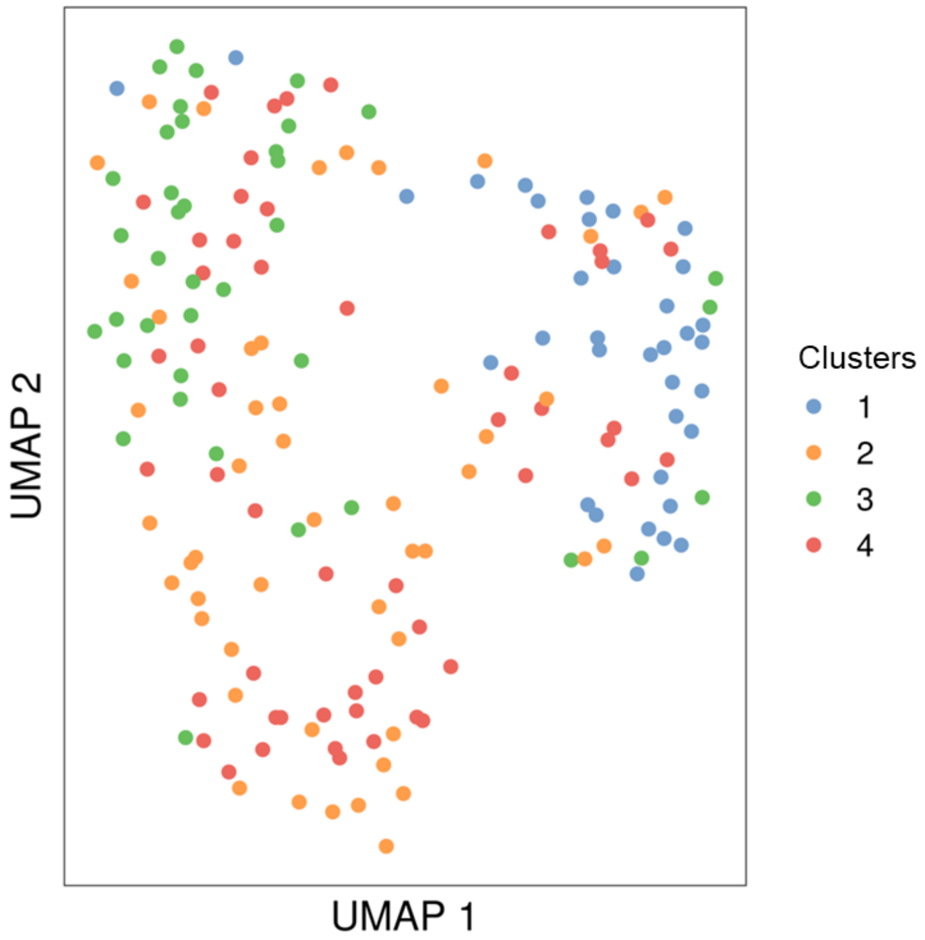


**Supplementary Figure S12. Visualizing cell states on deconvoluted CSE manifold in arthritis patients.**

UMAP plot of raw bulk RNA expression profile. Each dot represented a patient, and was colored according to the cell cluster inferred from the CSE of Monocytes.

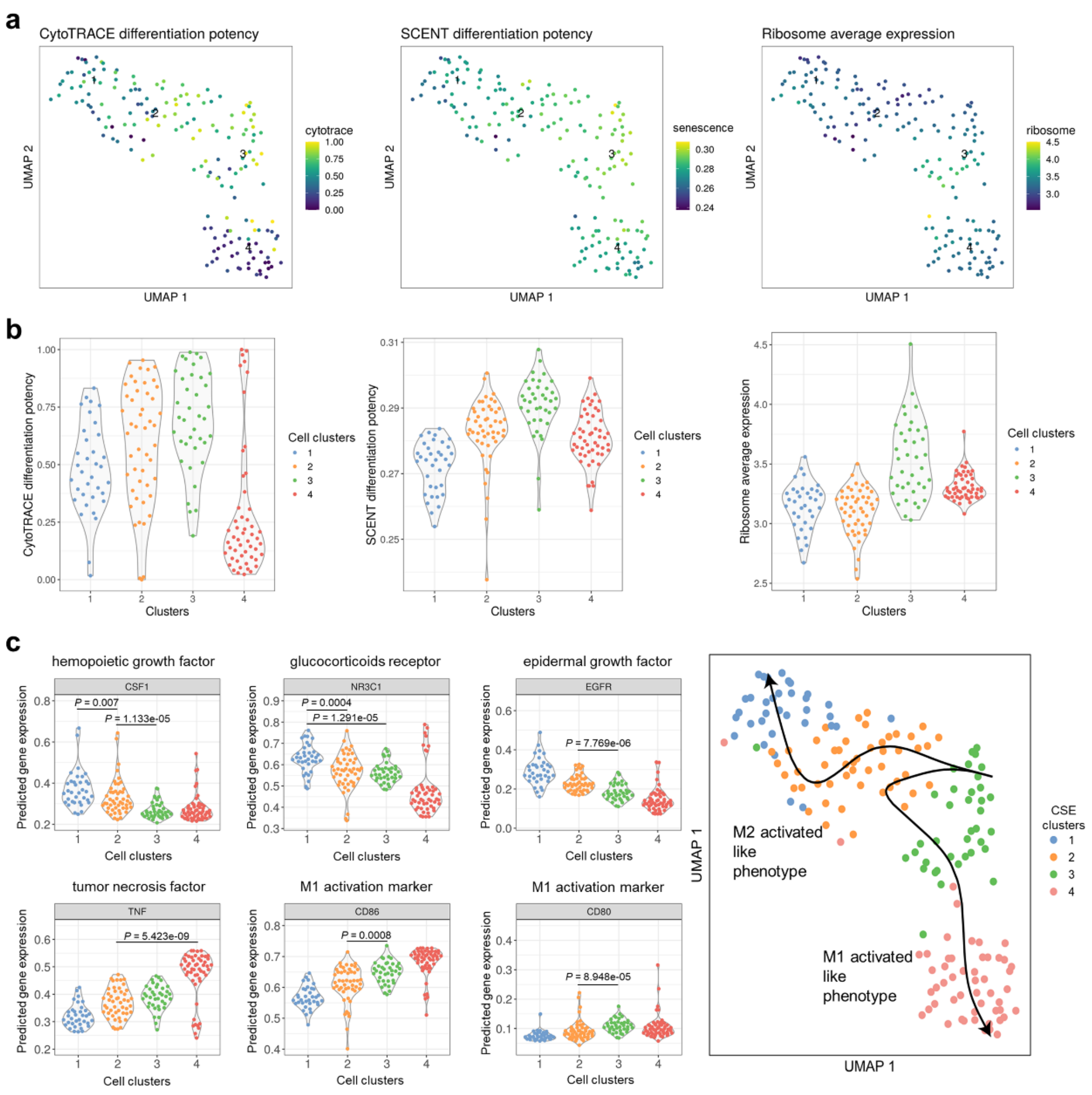


**Supplementary Figure S13. ENIGMA reveals pseudo-differentiation trajectory of monocytes.**

a. UMAP plots of CSE profile of Monocytes. Each dot represented a deconvoluted patient, and was colored according to CytoTRACE predicted differentiation potency score (left panel), SCENT predicted differentiation score (middle panel), and average expression of ribosome relation genes (right panel).
b. Violin plot of CytoTRACE predicted differentiation potency score (left panel), SCENT differentiation potency score (middle panel) and average expression of ribosome relation genes (right panel) in four clusters. Each dot was colored according to the cluster identity.
c. Violin plot of M1 macrophage and M2 macrophage related genes in four clusters (left panel). Right panel showed that UMAP plot of CSE profile of Monocytes. And each dot represented a deconvoluted sample/patient, and was colored according to the cluster identity. The principal curve was fitted through R package *Slingshot.*

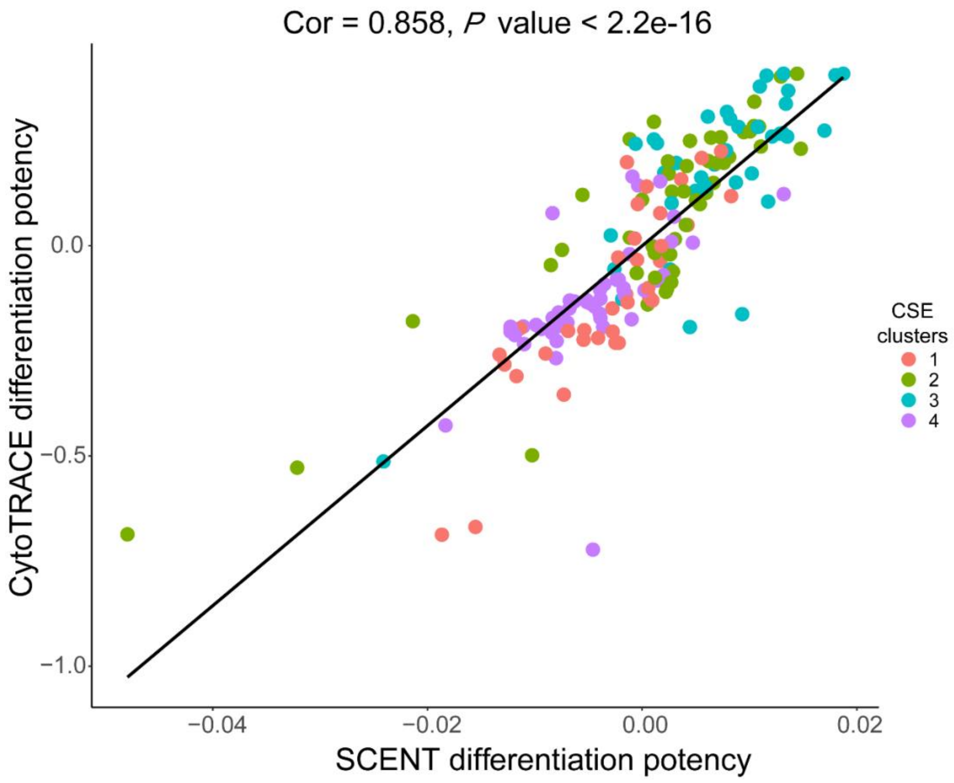


**Supplementary Figure S14. Concordance of differentiation potency prediction score on inferred monocyte expression profile.**

Scatter plot of CytoTRACE predicted differentiation potency (Y-axis) with SCENT predicted differentiation potency (X-axis) of predicted monocyte CSE profiles. Both differentiation potency scores were regressed out the variation of monocyte cell type fractions across samples. Each dot represented a sample and was colored according to the data clusters.

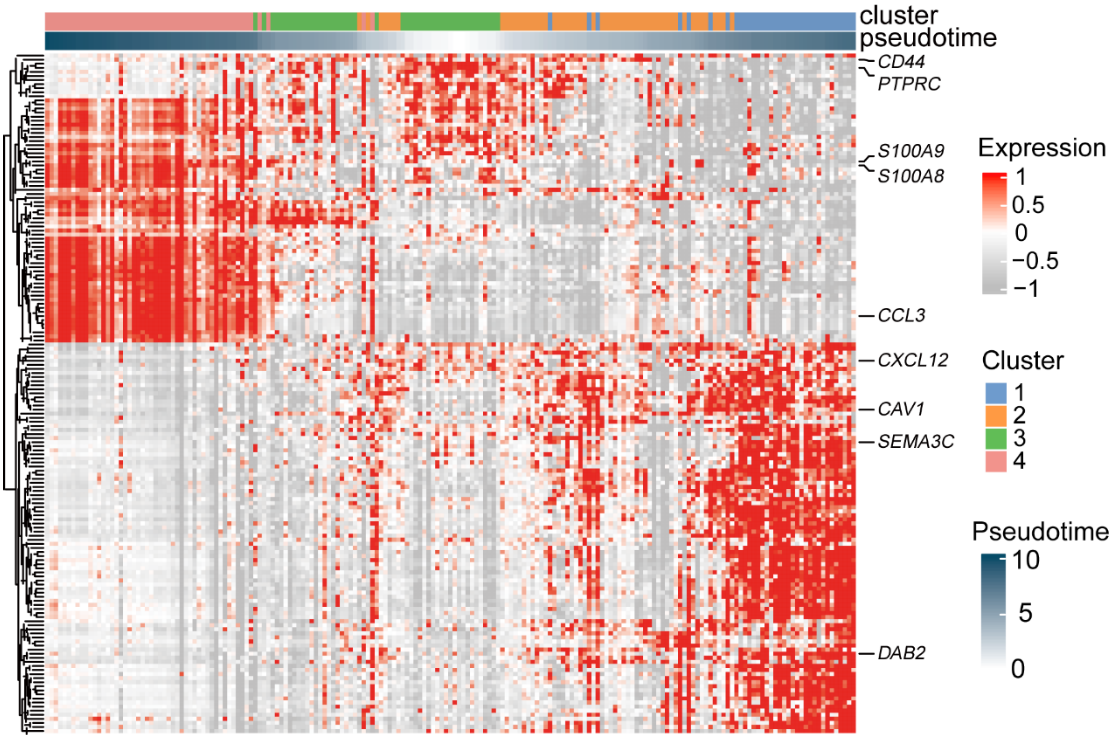


**Supplementary Figure S15. Heatmap of branch-specific genes**

Heatmap of branch-specific genes identified using the *tradeSeq* [5] through generalized additive model (*P*-value < 0.1) in Supplementary Figure S13c (right panel). Pseudo-time was calculated through R package *slingshot* [6]. The genes mentioned in the main text were shown at the right.

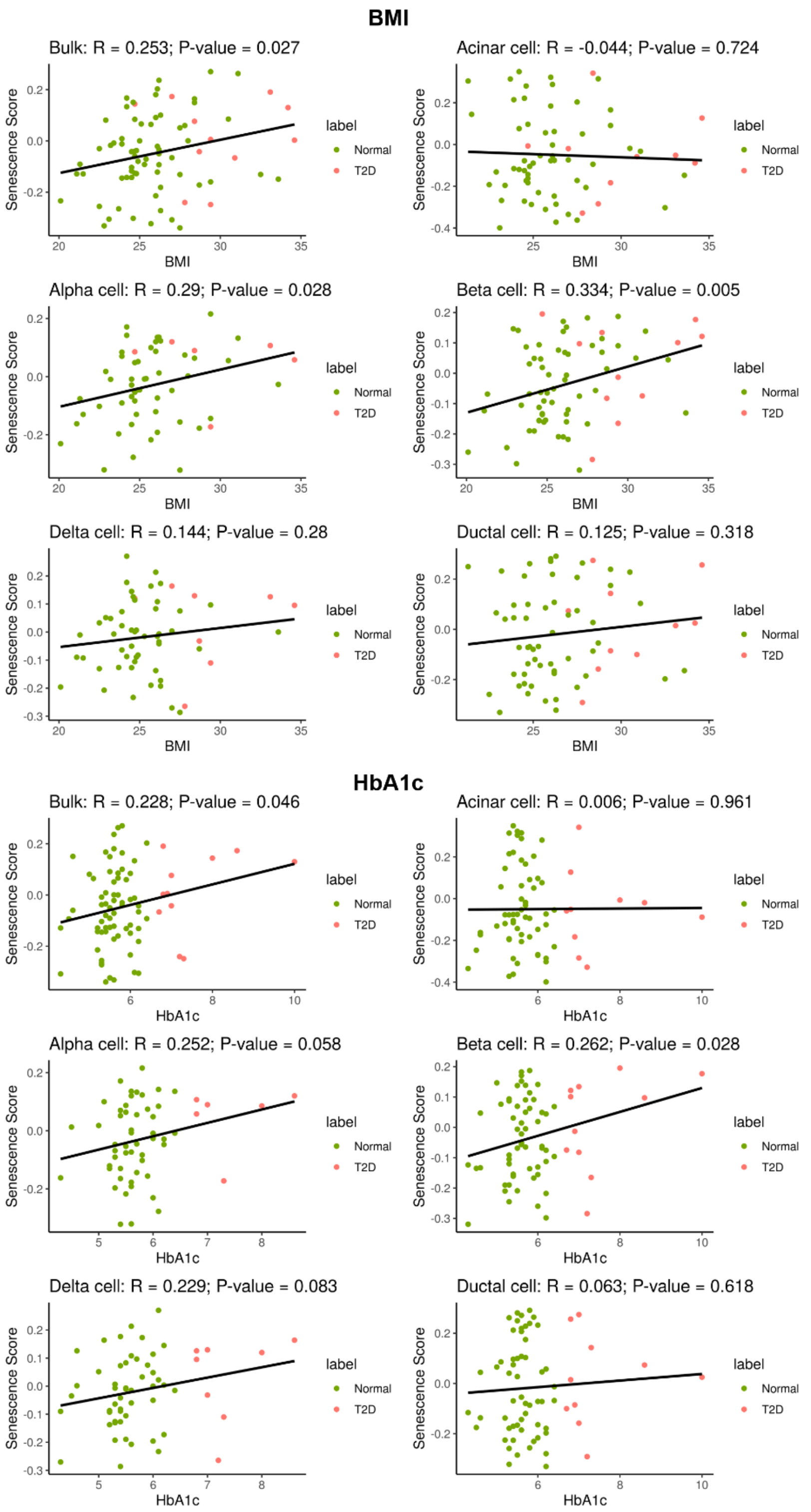


**Supplementary Figure S16. The correlation between cell type-specific senescence (CellAge) score with BMI and HbA1c levels in both normal and T2D donors.**

The scatter plots of senescence score calculated from expression profile (Y-axis) with the HbA1c level (X-axis). The correlation was calculated using Pearson correlation coefficients. Each dot represented a sample, and was colored according to the disease state (Normal or T2D). The senescence score was calculated based on the cell senescence signature provided by CellAge [7] database and gene set variation analysis tool named *GSVA* [8]

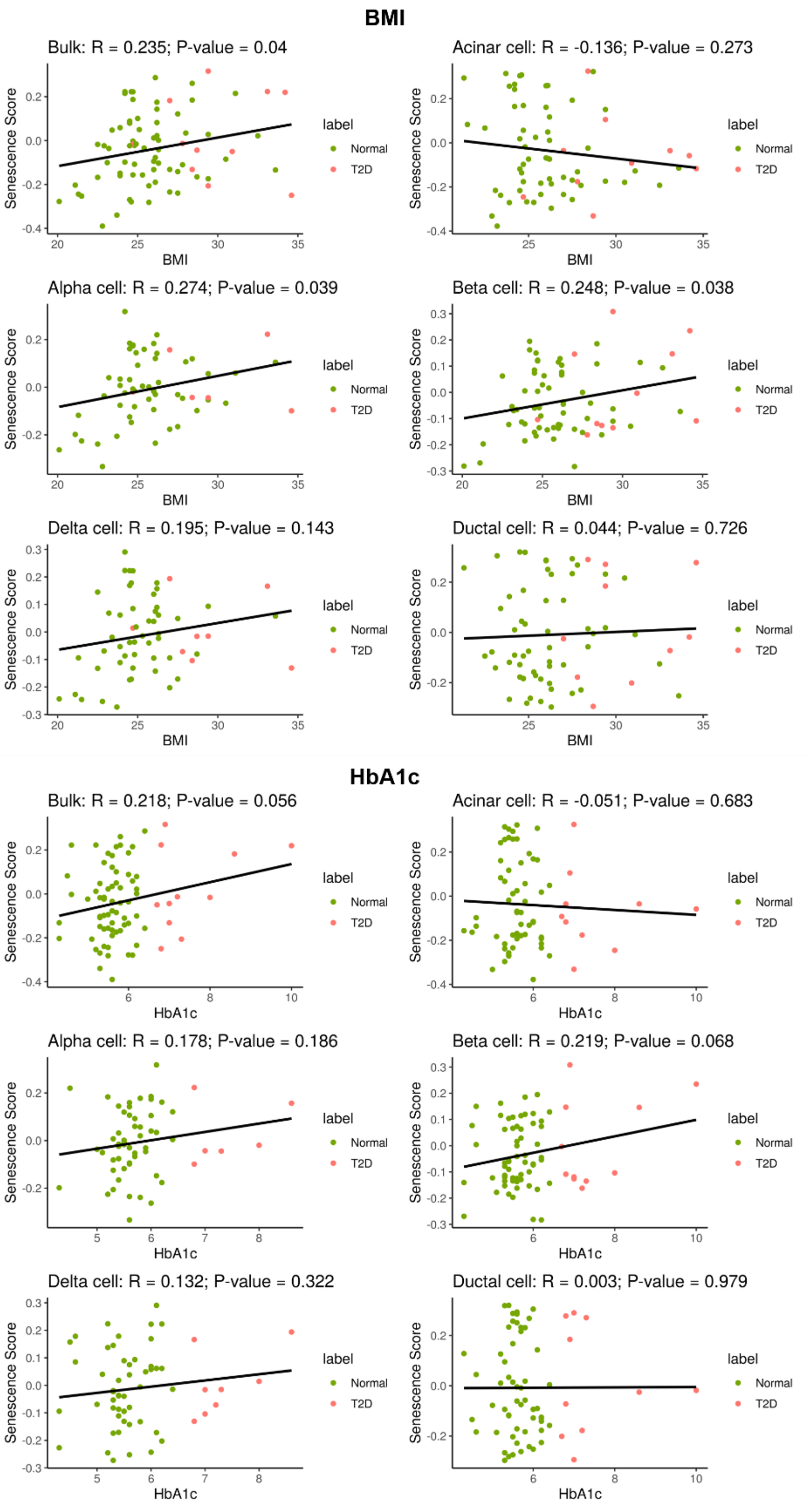


**Supplementary Figure S17. The correlation between cell type-specific senescence (SASP) score with BMI and HbA1c levels in both normal and T2D donors.**

The scatter plots of senescence score calculated from expression profile (Y-axis) with the HbA1c level (X-axis). The correlation was calculated using Pearson correlation coefficients. Each dot represented a sample, and was colored according to the disease state (Normal or T2D). The senescence score was calculated based on the cell senescence signature provided by CSGene [9] database and gene set variation analysis tool named *GSVA* [8].

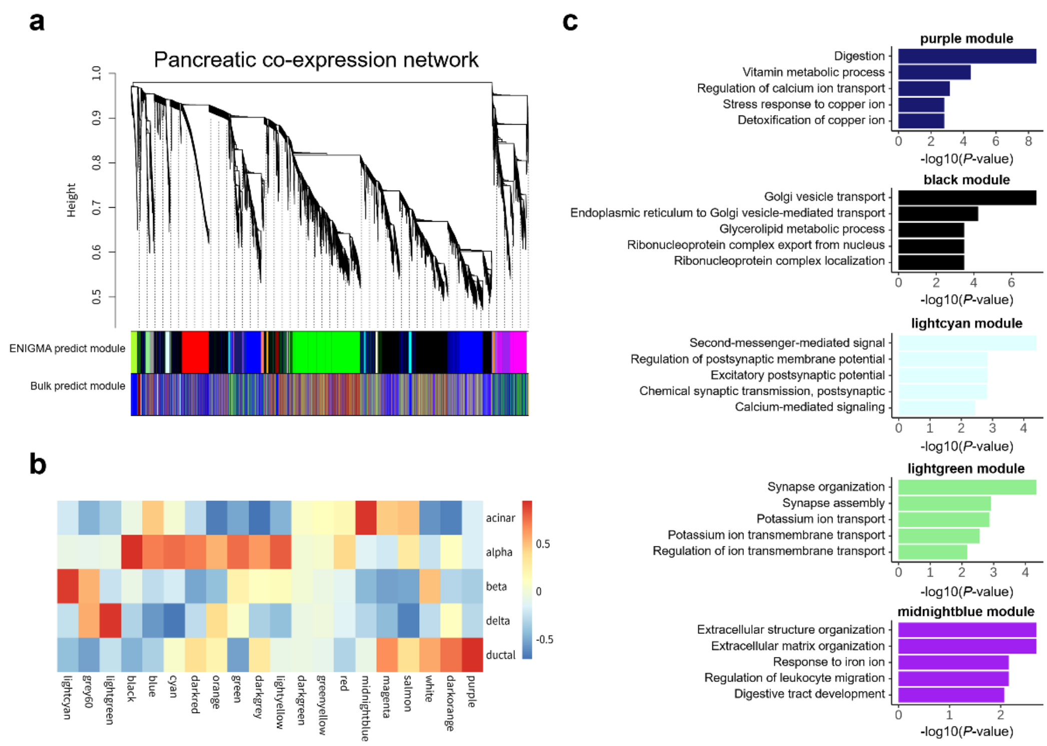


**Supplementary Figure S18. Cell type-specific gene co-expression network analysis in pancreas islet.**

a. Hierarchical cluster tree showing co-expression modules identified using WGCNA in all CSE profiles. Modules corresponding to branches were labelled by colors as indicated by the first color band underneath the tree. The modules identified by bulk RNA-seq were labelled by colors as indicated by the second color band underneath the tree.
b. Heatmap plot of correlation matrix between each module eigengene expression with the cell type index. Each entry was colored according to the correlation coefficients.
c. The top five most significant GO pathways enriched in each cell type that associated with modules. The pathways were ordered according to their significance, and the bar was colored according to its module names. GO analysis was conducted using R-package *clusterProfiler* [10].

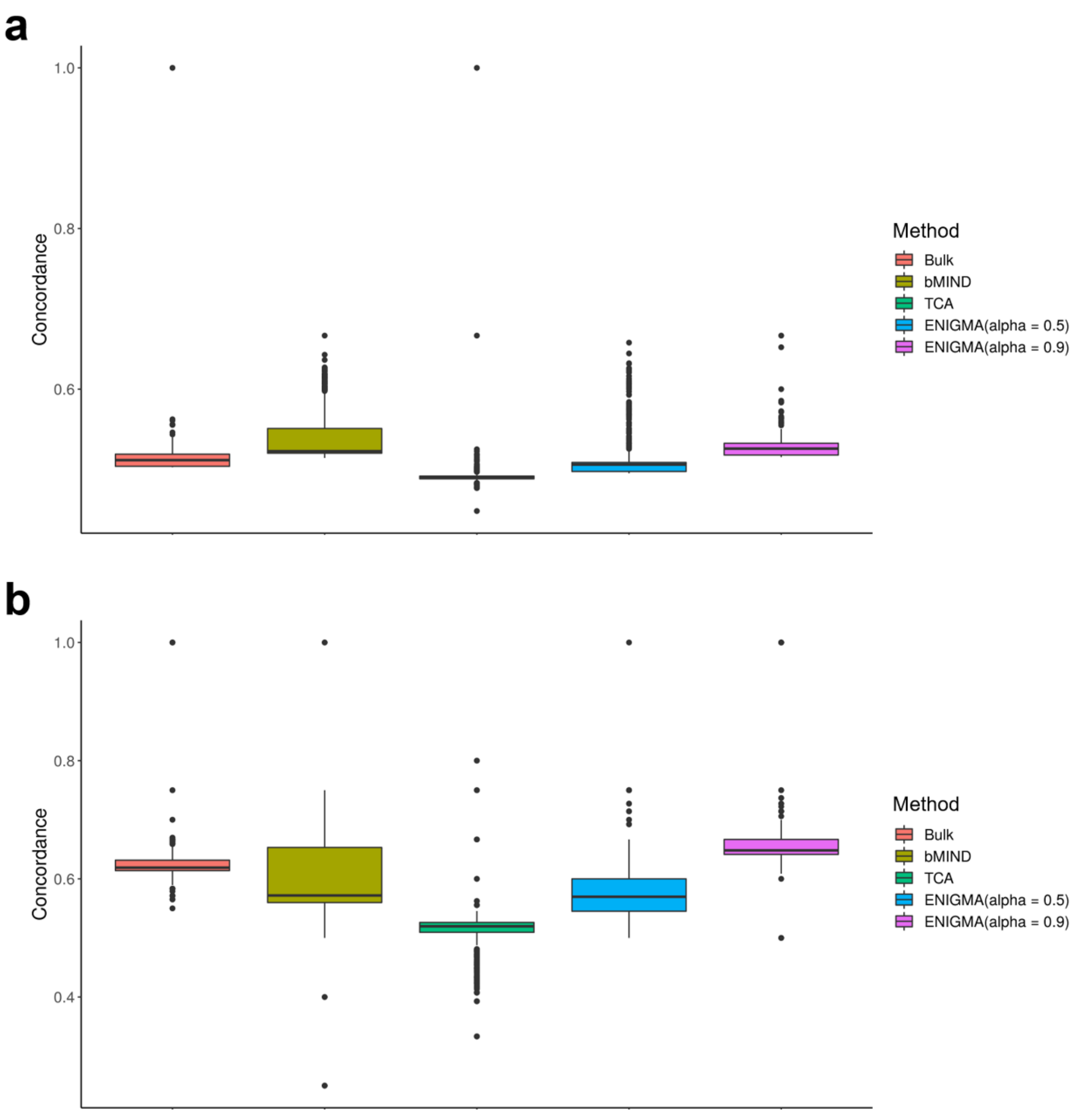


**Supplementary Figure S19. Optimize deconvolution models through improving beta cell-specific expression concordance with independent scRNA-seq datasets.**

The boxplots of gene differential expression concordance between beta cell specific expression profile inferred from TCA, bMIND, ENIGMA and independent pancreatic islet single cell RNA expression profile [11]. The phenotype was the disease state of the patients (Normal vs T2D). To make the comparison, the gene differential expression concordance between bulk RNA expression and pancreatic single cell RNA expression was also calculated. The gene differential expression concordance was calculated on all expressed genes (a) and top500 most differentially expressed genes (b). The reference matrix was constructed from pancreatic islet single cell RNA expression profile from a different study [12].

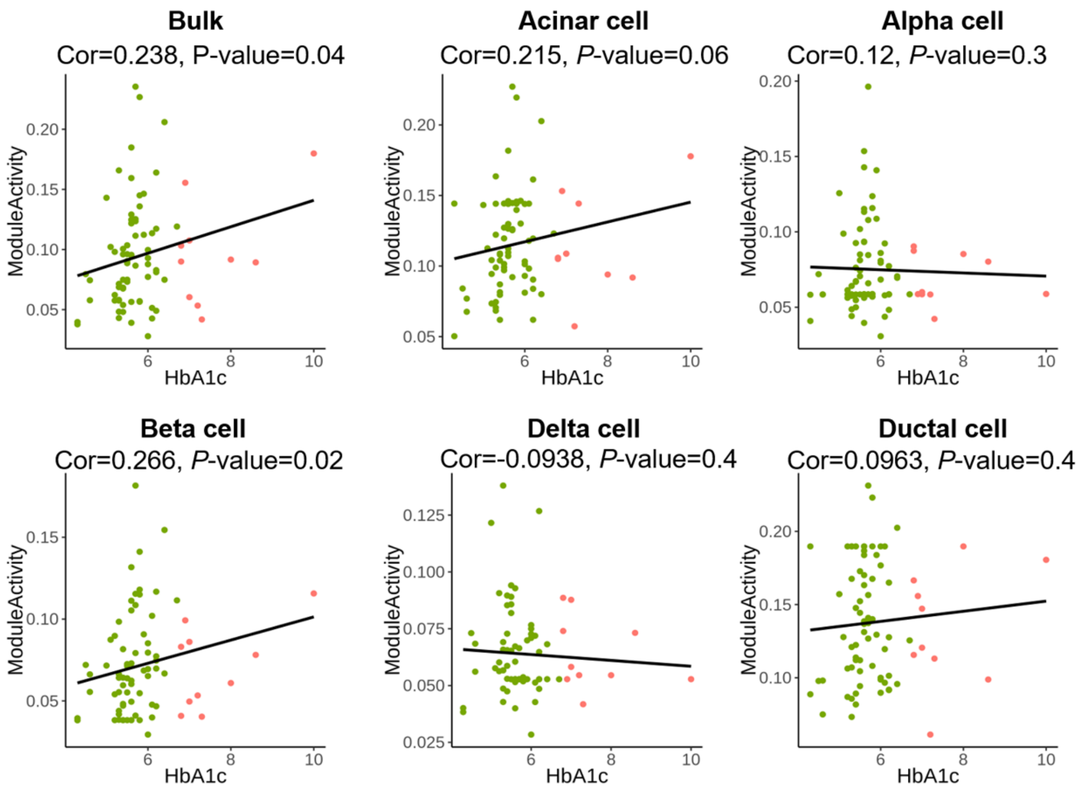


**Supplementary Figure S20. The correlation between apoptosis-associated module activity and HbA1c level for each cell type in both normal and T2D donors.**

The scatter plots of darkgreen module activity (Y-axis, Fig. 5d) with the HbA1c level (X-axis). The correlation was calculated using Spearman correlation coefficients (The module activity values were calculated through rank-based method, *AUCell* [13]). Each dot represented a sample, and was colored according to the disease state (Normal or T2D). The module activity was calculated through gene set variation analysis tool named *GSVA.*
